# Non-parametric GWAS: Another View on Genome-wide Association Study

**DOI:** 10.1101/2022.11.11.516099

**Authors:** Xiaoyue Hu, Shizhou Yu, Hangjin Jiang

## Abstract

Genome-wide association study (GWAS) is a fundamental step for understanding the genetic link to traits (phenotypes) of interest, such as disease, BMI and height. Typically, GWAS estimates the effect of SNP on the phenotype using a linear model by coding SNP as working code, {0, 1, 2}, according to the minor allele frequency. Looking inside the linear model, we find that the coding strategy of SNP plays a key role in detecting SNPs contributed to the phenotype. Specifically, a partial mismatch between the order of the working code and that of the underlying true code will lead to false negatives, which has been ignored for a long time. Motivated by this phenomenon, we propose an indicator of possible false negatives and several non-parametric GWAS methods independent of coding strategy. Results from both simulations and real data analysis show the advantages of new methods in identifying significant loci, indicating their important complementary role in GWAS.

## 1 Background

With the innovation of high throughput genotyping technologies of single nueleotide polymorphism (SNP), genome-wide association studies (GWAS) have witnessed considerable progress in the identification of significant genetic factors. Since the first GWAS published, more than 50,000 associations have been reported between SNPs and diverse traits or diseases of humans, animals, and plants (Tam et al., 2019). GWAS has been widely used for its high efficacy in scenarios of disease susceptibility (Bouaziz et al., 2011), disease biomarkers identification (Bossé and Amos, 2018), risk prediction (Ji et al., 2021), and therapy optimization (Prokop et al., 2018).

The most common analysis in GWAS is single marker regression, putting SNPs as independent variables and phenotypes as dependent variables to identify significant loci (Dehghan, 2018). Given the nature of phenotypes and the purpose of GWAS, a linear, logistic, or Cox regression model is used to evaluate the impact of SNP on the phenotype. However, the majority of SNPs labor in non-coding regions, making it difficult to understand the biological mechanism of how genetic factors influence phenotypes (Giambartolomei et al., 2018). One popular approach to facilitate the understanding is to integrate molecular traits as mediators, such as gene expression (eQTL), DNA methylation (mQTL), and histone modification levels (haQTL), collectively known as xQTL (Ng et al., 2017). By jointly analyzing the results of GWAS and xQTL, researchers have found that variants, which drive the associations in GWAS and affect xQTL expressions at the same time, shed light on the machinery of gene regulation (Fromer et al., 2016; Hannon et al., 2017).

In line with a pattern of polygenic inheritance, more and more SNPs have been identified by performing GWAS with an increasing sample size or meta-analyses of summary statistics from a surging number of studies (Yang et al., 2012). However, the need of large sample size is a primary limitation, since it is costly to collect or not feasible for many species, such as animals and plants (Tam et al., 2019). Therefore, new approaches are strongly in need to achieve higher power than GWAS with the same sample size, so as to pinpoint more significant loci. To facilitate numerical analysis, SNPs are typically coded as {0, 1, 2} according to allele frequency, say, major allele as 0, heterozygote as 1, and minor allele as 2, and then a linear model is used to evaluate the effect of SNP on the phenotype. There are two major concerns about this approach. First, it assumes that SNPs follow an additive model. However, this may not be held under many circumstances. Although additive model has reasonable power to detect both additive and dominant effects, it may be underpowered to detect SNPs following recessive model or other genetic mode (Bush and Moore, 2012). Second, the coding strategy of SNPs implicitly introduces some numerical assumptions to SNPs, and may lead to a low power in some cases, which has been ignored for a long time. In this paper, we focus on this concern to consider the impact of coding strategy of SNPs on the power of GWAS and propose alternative methods to increase the power.

To understand when and how the coding strategy leads to power loss, we analyze the classical linear model theoretically and find that a partial mismatch between the order of working code and that of the underlying true code will result in possible false negatives. Thus, we propose non-parametric methods that do not depend on the coding strategy of SNPs for both continuous and binary traits. Simulations are given to evaluate the performance of different methods under different cases, and to better understand our theoretical analysis. After that, we illustrate the benefits of non-parametric GWAS in seven real datasets covering human, plants, and animals. In the analysis, we identify many significant SNPs and genes neglected by GWAS, and some of these loci were verified to have strong association with the corresponding traits in the literature. Importantly, we discover that different methods have diverse performance on the same dataset, which might confuse users. Along this line, discussion on how to select statistical methods for GWAS is taken to make clear the confusion.

## 2 Results

### 2.1 GWAS based on linear model may have false negatives

Typically, GWAS for finding SNPs related to the trait *Y* is to test whether *β* = 0 or not in the following linear model between SNP (*G*) and *Y*:

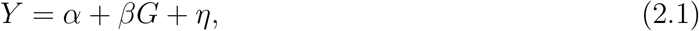

where *α* is the background effect, *β* is the genetic effect, and *η* is the noise term assumed to be normal with mean 0.

Taking the working code for *G* as *c_i_* = *i* for *i* = 0,1, 2, *β* is estimated as,

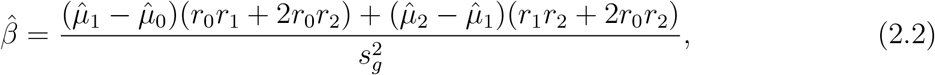

where 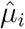 is the conditional sample means of *Y* given *G* = *c_i_*, *r_i_* = *n_i_/n* is the ratio of sample size for *G* = *c_i_* over the total sample size *n*, and 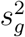 is the sample variance of *G*.

Now, we define the true code as the code giving 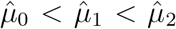. That is, the genotype coded as “0” has weakest effect on the phenotype, and that coded as “2” has the strongest effect. This is the basic assumption underlying GWAS. However, the working code may be different from the true code and gives different order of 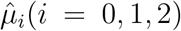 (see Table 1), which also can be explained by a general model (2.3) with details given in Methods. This phenomenon may introduce a critical problem to the estimation of *β*. Generally, according to equation (2.2), if the working code gives 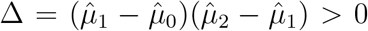, then the effect of SNP on the trait is not affected by the working code. Otherwise, we may have false negatives due to the coding strategy of SNP. In detail, when Δ < 0, which means the information from the group mean difference may cancel out and lead to 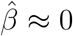 even if the true β is not 0, and thus reports a false negative. Typically, when *n*_1_ = *n*_2_ = *n*_3_, 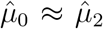 gives 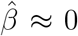, despite of 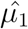. Thus, Δ < 0 is a signal for possible false negatives. Results in Simulations confirm this statement, further evidenced by results in Real Data Analysis. In summary, when the order of working code and that of true code are partially mismatched, GWAS may have a lower power to detect meaningful SNPs.

**Table 1:** Possible order of 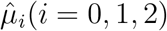 with working code *c_i_* = *i*(*i* = 0,1, 2).

|  |  |  |
| --- | --- | --- |
| C1: $\hat{\mu}_0 < \hat{\mu}_1 < \hat{\mu}_2$ | C2: $\hat{\mu}_1 < \hat{\mu}_0 < \hat{\mu}_2$ | C3: $\hat{\mu}_2 < \hat{\mu}_0 < \hat{\mu}_1$ |
| C4: $\hat{\mu}_0 < \hat{\mu}_2 < \hat{\mu}_1$ | C5: $\hat{\mu}_1 < \hat{\mu}_2 < \hat{\mu}_0$ | C6: $\hat{\mu}_2 < \hat{\mu}_1 < \hat{\mu}_0$ |

Motivated by these facts, we introduce non-parametric GWAS by considering the following general model:

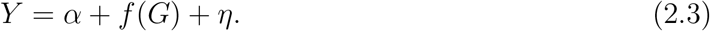

That means, the genetic effect is not directly put on the trait *Y*, but through an unknown function *f*. The linear model is a special case of this general model by taking *f*(*x*) = *βx*. The biological implication is that the genetic effect of each SNP is unknown in practice.

To get rid of the coding strategy of SNPs, we propose several non-parametric GWAS methods: W-GWAS, Ca-GWAS, Chi-GWAS, KS-GWAS, Co-GWAS, and HC-GWAS for continuous phenotypes and Chi2-GWAS for binary traits. A brief description of these methods is shown in Table 2 with details provided in Methods. In summary, W-GWAS and Ca-GWAS both utilize pairwise Wilcoxon rank-sum test, and the only difference is that the p-value from W-GWAS is Bonferroni corrected while p-value in Ca-GWAS is a combined p-value of p-values from pairwise comparison. Chi-GWAS uses the the sum of squared differences of group means, whose limit distribution follows a Chi-squared distribution. Just as its name, KS-GWAS makes use of pairwise KS test. Co-GWAS is based on the comparison between conditional distributions to test whether these distributions under different SNPs are the same or not. Finally, HC-GWAS integrates all the former methods by p-value combination, following the idea proposed by Li et al. (2022). For the binary scenario, Chi2-GWAS is based on the adjusted Chi-squared test.

**Table 2:** Description of non-parametric GWAS methods

| Type | Method | Description |
| --- | --- | --- |
| Continuous | W-GWAS | based on pairwise mean test with Bonferroni adjusted p-value |
|  | Ca-GWAS | based on pairwise mean test with p-value<br>combined by Cauchy transformation |
|  | Chi-GWAS | sum of squared differences of group means |
|  | KS-GWAS | pairwise KS test with adjusted p-values |
|  | Co-GWAS | based on comparison between conditional distributions |
|  | HC-GWAS | integrating five methods by p-value combination |
| Binary | Chi2-GWAS | based on adjusted Chi-squared test |

### 2.2 Simulations

In this section, simulations are taken to evaluate the power of non-parametric GWAS methods and compare them with GWAS. When the phenotype is continuous, we compare six nonparametric GWAS methods (W-GWAS, Ca-GWAS, Chi-GWAS, KS-GWAS, Co-GWAS, and HC-GWAS) with three different versions of GWAS: (1) GWAS with default code {0, 1, 2} given according to minor allele frequency; (2) GWAS with working code having the same order of the true genetic code, called GWAS1; and (3) GWAS with true code, called GWAS2. The design of these different versions is to evaluate the impact of coding strategy on the power of GWAS. Similarly, for binary phenotypes, Chi2-GWAS is also compared with GWAS, GWAS1, and GWAS2.

The aim of simulation is to (a) provide numerical evidence on the possibility of false negatives from GWAS and (b) show advantages and disadvantages of different methods. According to the analysis given before and in Methods, we focus on three different situations: (S1) the order of working code is the same as that of true code (Example A); (S2) these two orders are partially mismatched (Examples B and C); and (S3) they are fully unmatched (Example D); Details of these simulation examples are given in Methods. The power of different methods is estimated from 200 independent repetitions under significance level 0.05. Simulation results are presented in Figure 1 (examples with continuous phenotypes) and Figure 2 (examples with binary phenotypes).

**Figure 1:**
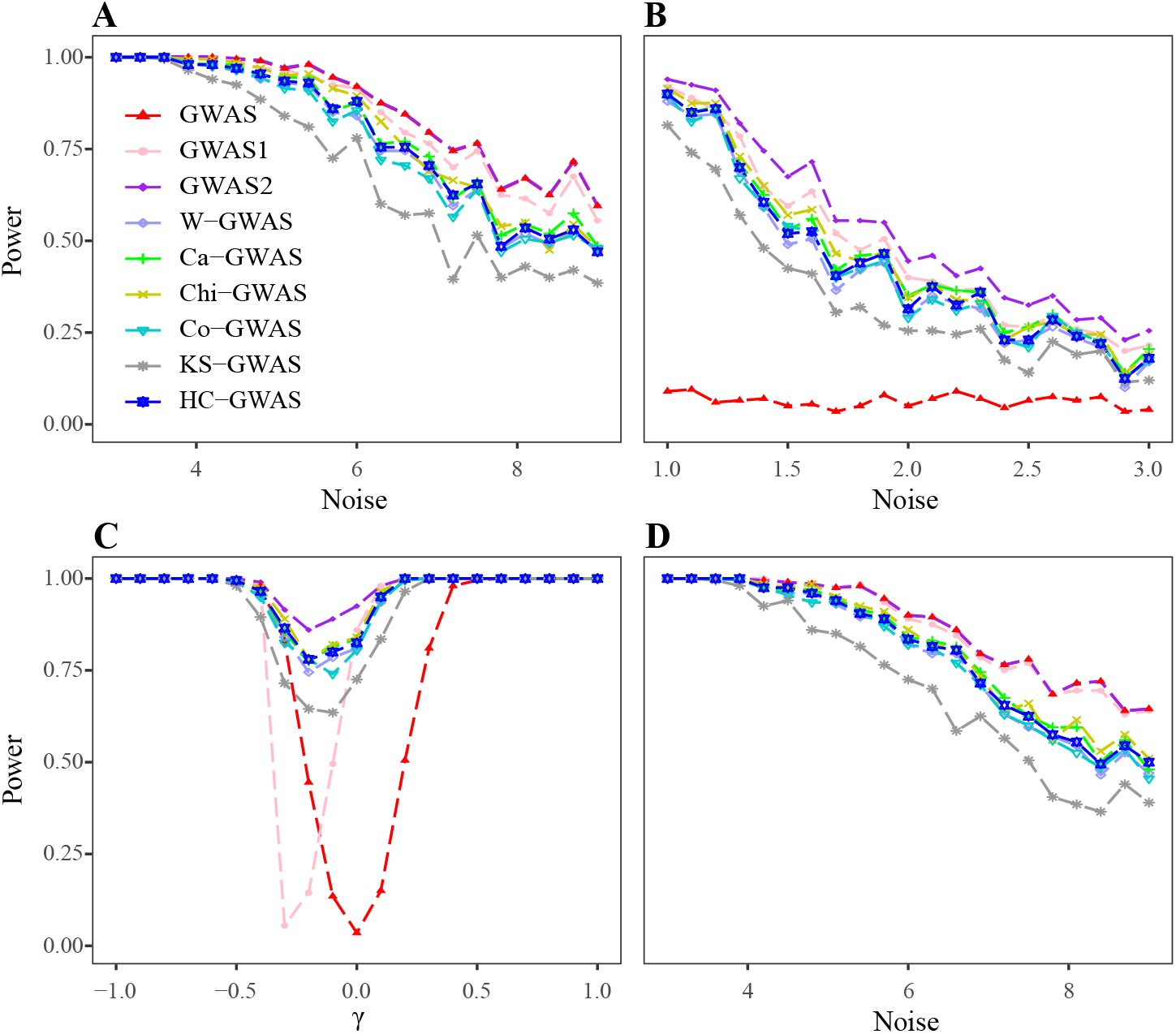
Power of different methods in continuous examples. **A**) Case S1 that the order of working code is the same as that of true code; **B-C**) Case S2 that these two orders are partially mismatched; **D**) Case S3 that they are fully unmatched. The results show that if we know exactly the true genetic code, GWAS is the best method to use. However, it is usually unknown to us in real problems. Thus, we suggest to use non-parametric methods as a complement to the traditional GWAS to avoid high false negatives.

**Figure 2:**
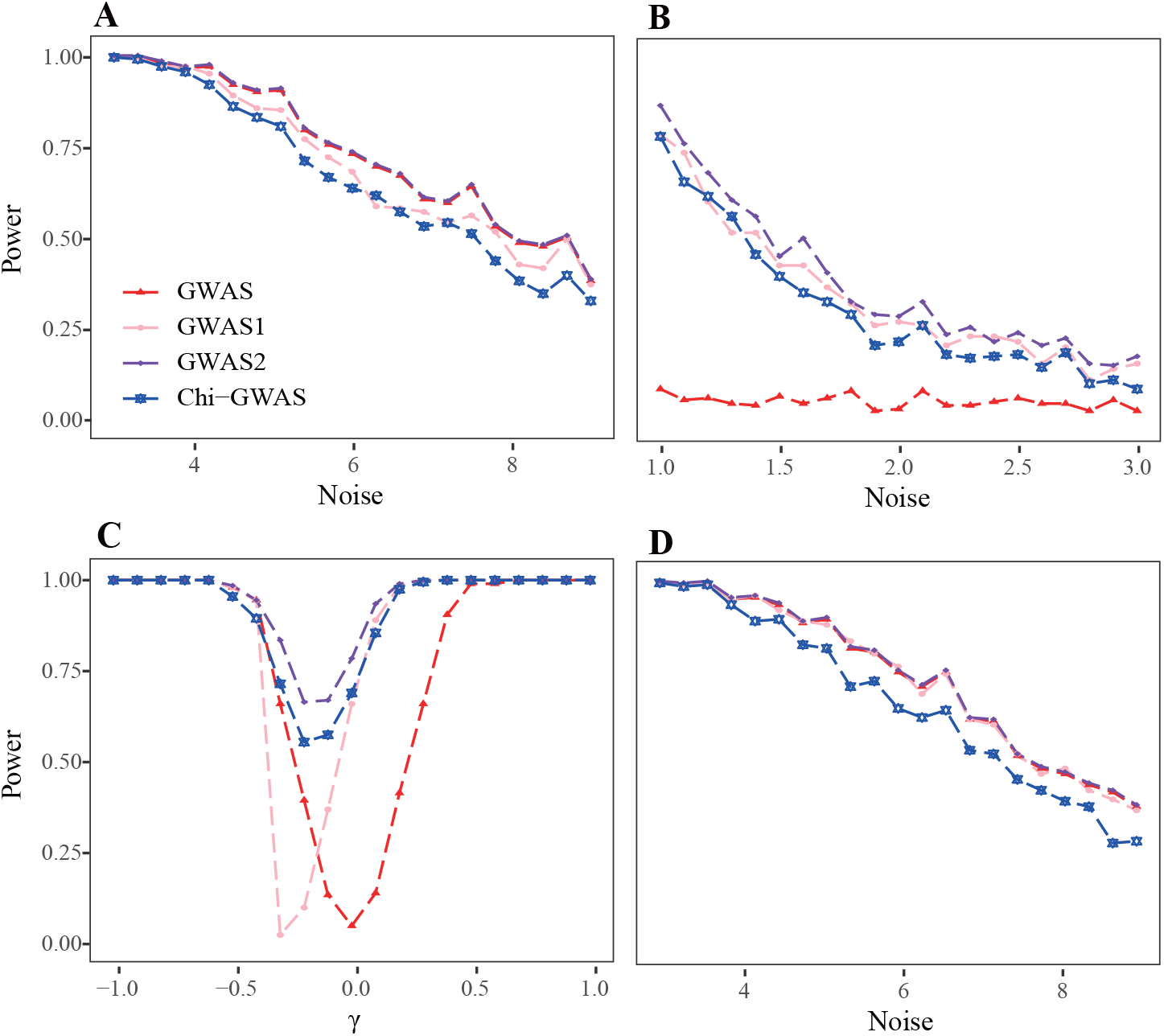
Power of different methods in binary examples. **A**) Case S1 that the order of working code is the same as that of true code; **B-C**) Case S2 that these two orders are partially mismatched; **D**) Case S3 that they are fully unmatched. The results are in line with that of continuous examples.

For both cases of phenotypes, GWAS2, GWAS with true genetic code, performs the best among these methods. When the order of two codes are partially matched, as in Example B and C, GWAS, with default working code {0,1,2}, always has the worst performance. It suggests that traditional GWAS has a low power to identify true loci whose genetic mode do not follow the default additive model. Moreover, GWAS1, whose working code has the same order as the true genetic code, sometimes has a comparable performance with GWAS2, as shown in Example A, B, and D. Nevertheless, when *γ* in Example C is close to −0.3, GWAS1 gives a very low power which even drops below 0.1. Note that the behavior of GWAS and GWAS1 are different in Example C. GWAS has the lowest power when *γ* ≈ 0, corresponding to the case 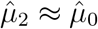 leading to 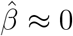. Differently, GWAS1 has the lowest power when *γ* ≈ −0.3 and the information of 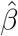 only comes from 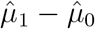, which is also small in this example, and thus leads to a lower power compared with other methods. Finally, the behavior of GWAS2 and non-parametric GWAS in this example are similar, as they all have a relatively lower power when *γ* ≈ —0.3. The result illustrates that if we have a good guess on the genetic mode, we may gain some power sometimes but may also have a lower power at other times. All these findings are consistent with our theoretical analysis.

Non-parametric methods demonstrate quite high power in all examples, showing their stability in identifying correlated SNPs with genetic mode in either order. Especially, they outperform GWAS in cases with partial mismatch (Examples B and C for both types of phenotype). Among these methods, W-GWAS and Ca-GWAS have comparable performance as their underlying algorithms are similar. HC-GWAS behaves normally without extreme conditions and its power is in the medium. Therefore, HC-GWAS could be deemed as a good representative for all the other non-parametric methods. In the binary scenario, the performance of Chi2-GWAS is in accordance with that of non-parametric GWAS methods in the continuous examples.

In summary, consistent with our theoretical analysis, the coding strategy is important to the performance of GWAS. When the order of working code and that of true code are partially mismatched, the performance of GWAS is disappointed. However, even if we set SNPs in the correct order, we may also have a lower power under certain circumstances, not to mention that we do not know the true genetic code in real problems. Nevertheless, we still have to admit that GWAS outperforms non-parametric methods in some cases. Accordingly, we suggest to use non-parametric methods as a complement to the traditional GWAS to avoid high false negatives and detect SNPs truly contributed to the response. According to our results, HC-GWAS is recommended as a summary statistic of other 5 non-parametric methods for continuous phenotypes.

### 2.3 Real Data Analysis

To explore in reality the aforementioned problem that GWAS might lose power in SNPs with partial mismatch, we compare the performance of GWAS and non-parametric GWAS in seven real datasets of different species including cucumber, cotton, sheep, pig, mouse, chicken, and human. Table 3 gives summary information of them with details provided in Methods. For continuous phenotypes, we utilize GWAS and 6 non-parametric methods to identify significant SNPs and genes. For binary traits, GWAS and Chi2-GWAS are implemented. Furthermore, pathway enrichment analysis is conducted if possible.

**Table 3:**
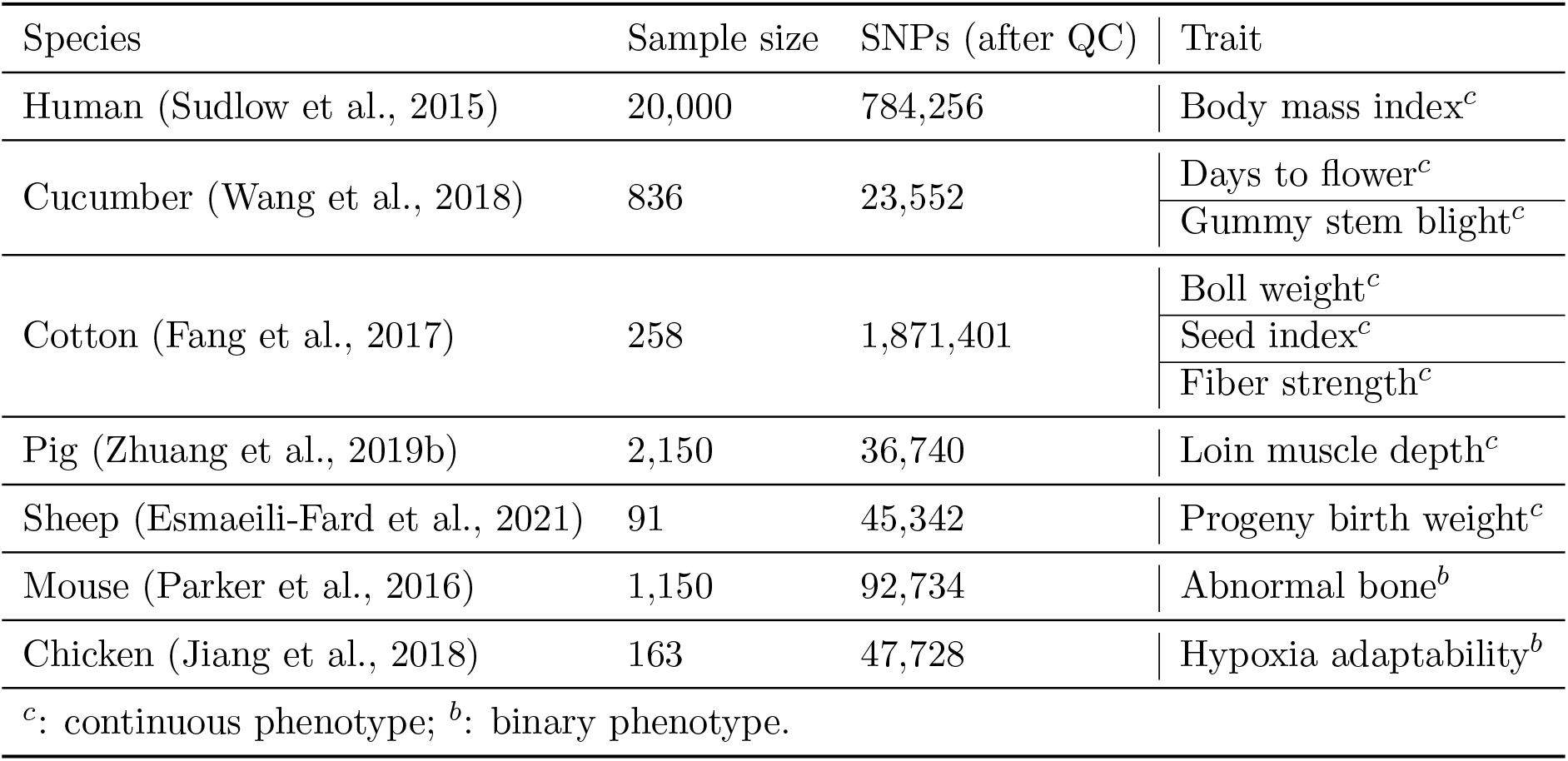
Summary information for datasets used in this study.

#### 2.3.1 Real data examples provide empirical evidence on the drawback of GWAS

GWAS and all the non-parametric methods are used to identify significant SNPs for different traits in different datasets (Table 3). Figure 3 shows a general relationship between the numbers of significant SNPs detected by GWAS and non-parametric GWAS in these datasets. More detailed relationship is shown in Figure 4 and Figure S1. Generally, GWAS may identify the most significant SNPs but neglect some loci deemed to be significant by other methods. Also, there are cases when GWAS only detects a small part of the significant SNPs. To provide evidence on the potential importance of SNPs detected by different methods, Table 4 shows part of the significant SNPs reported to be correlated with the trait of interest in literature but only detected by GWAS or one of these non-parametric methods.

**Figure 3:**
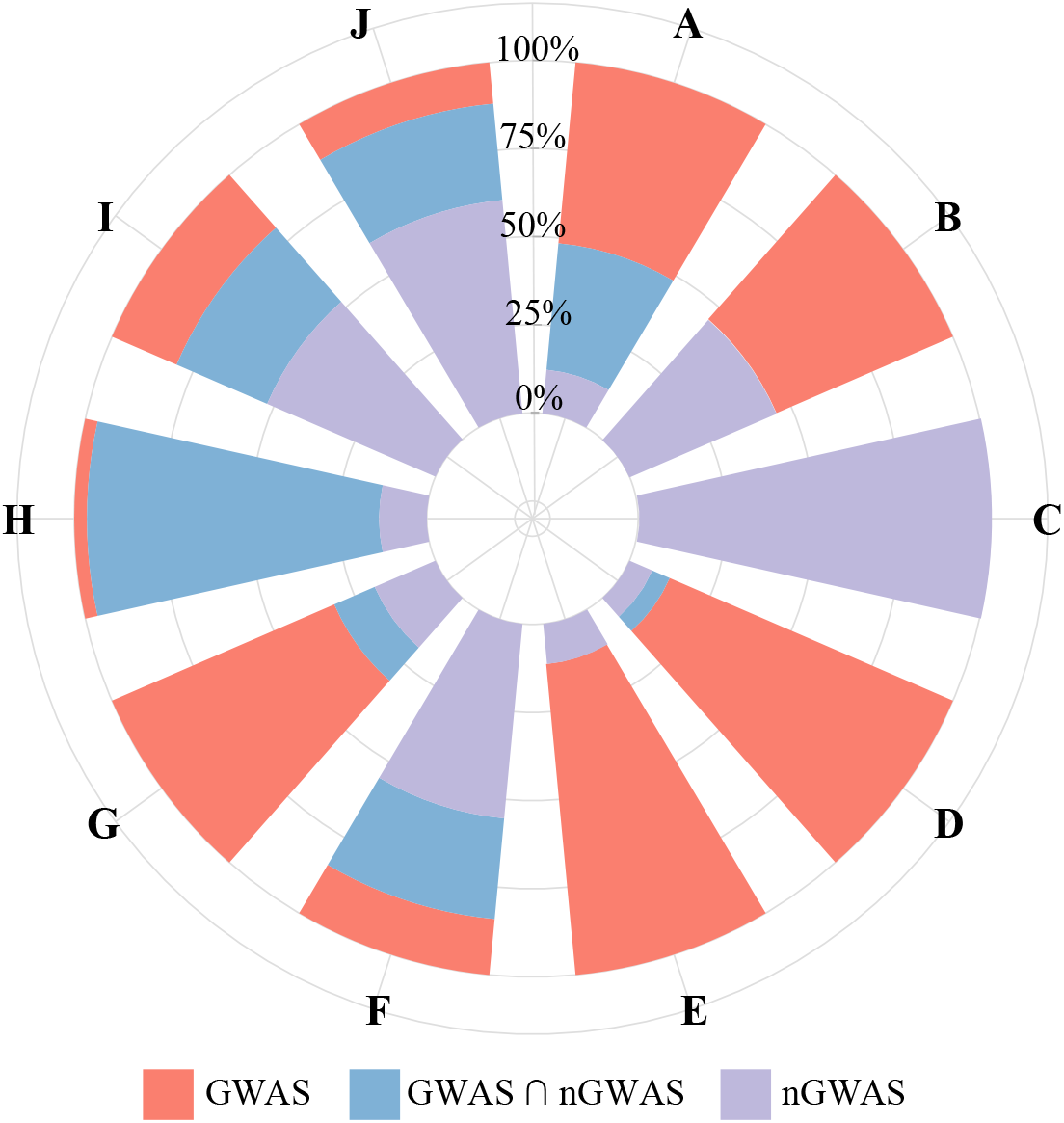
Percentage of significant SNPs detected by GWAS and non-parametric GWAS (nGWAS) in different datasets. **A**) Days to flower of cucumber; **B**) Gummy stem blight of cucumber; **C**) Boll weight of cotton; **D**) Seed index of cotton; **E**) Fiber strength of cotton; **F**) Progeny birth weight of sheep; **G**) Loin muscle depth of pig; **H**) Abnormal bone of Carworth Farms White (CFW) mouse; **I**) Hypoxia adaptability of chicken; **J**) Body mass index of human.

**Figure 4:**
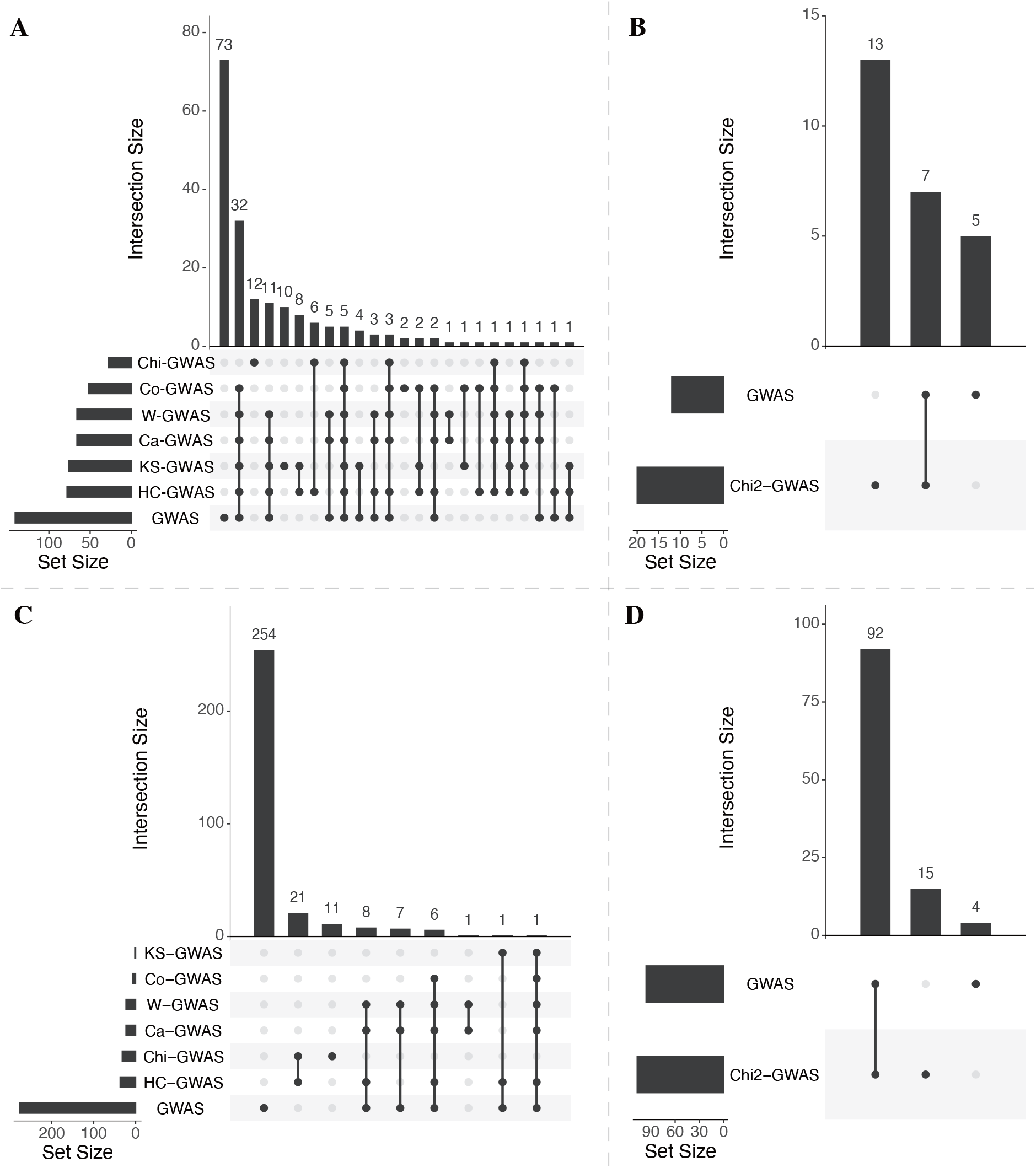
Relationship between significant SNPs detected by different methods in four datasets. **A**) Days to flower of cucumber; **B**) Hypoxia adaptability of chicken; **C**) Seed index of cotton; D) Abnormal bone of Carworth Farms White (CFW) mouse.

**Table 4:**
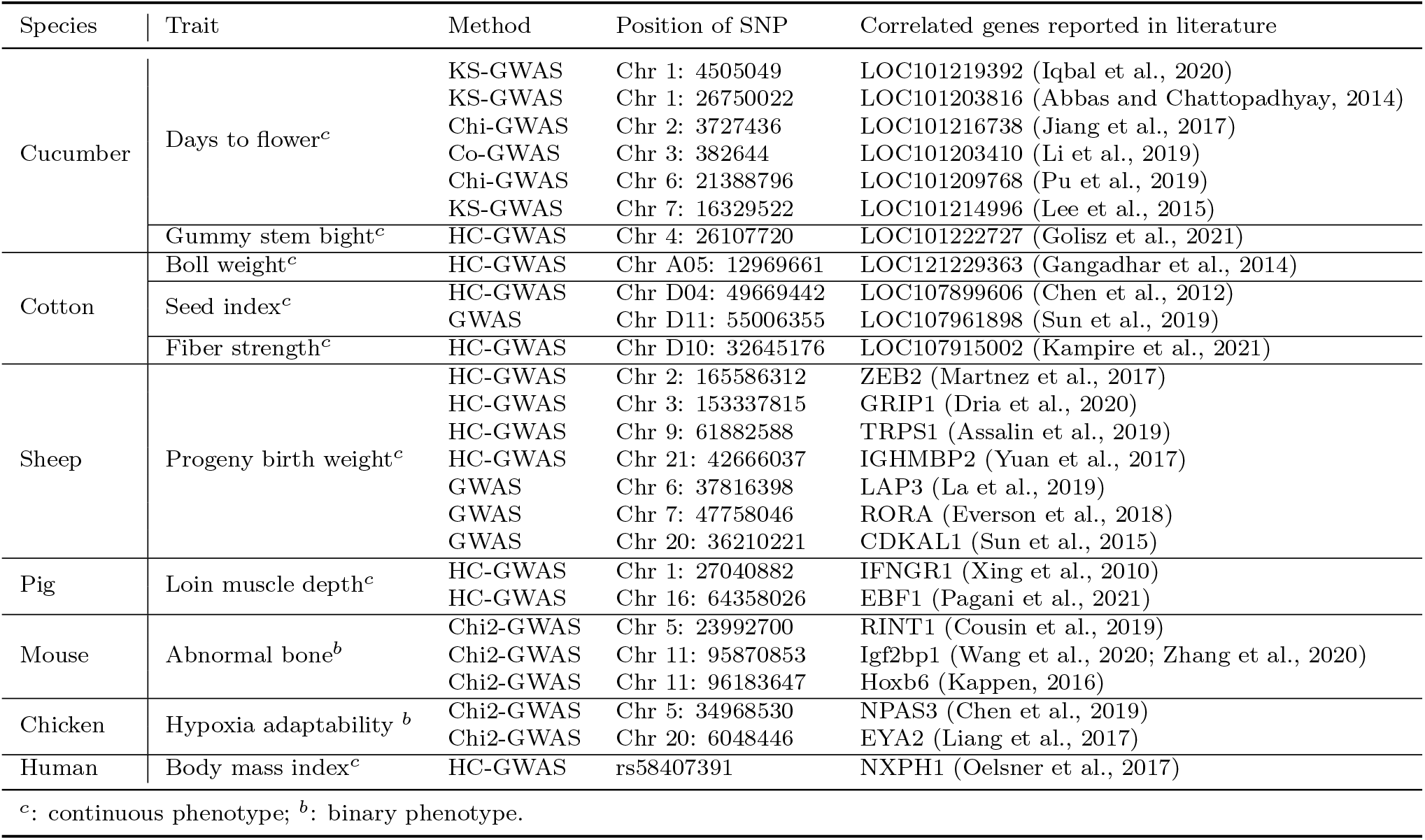
Informtion of significant SNPs detected by only one method

It is important to seek for the reason why GWAS misses so many SNPs. Given the working code {0, 1,2}, the empirical conditional means of phenotype are derived from 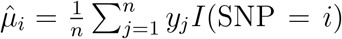 (SNP = *i*), for *i* = 0, 1, 2, where *n* is the sample size and *I*(*A*) is the indicator function taking value 1 when *A* is true, otherwise 0. As shown in Table 1, there are 6 possible orders between 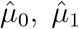 and 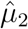, say C1-C6. Among these orders, C1 and C6 give 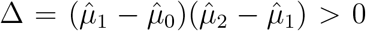, and C2-C5 give Δ < 0. To verify our analysis that GWAS is more likely to lose SNPs with Δ < 0, we classify the significant SNPs detected (or ignored) by GWAS into 6 classes according to the order of corresponding 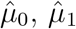 and 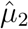, and get the distribution of SNP numbers in each class. Results in Figure 5 and Figure S2 show that GWAS tends to miss most of significant SNPs with Δ < 0, consistent with our theoretical analysis.

**Figure 5:**
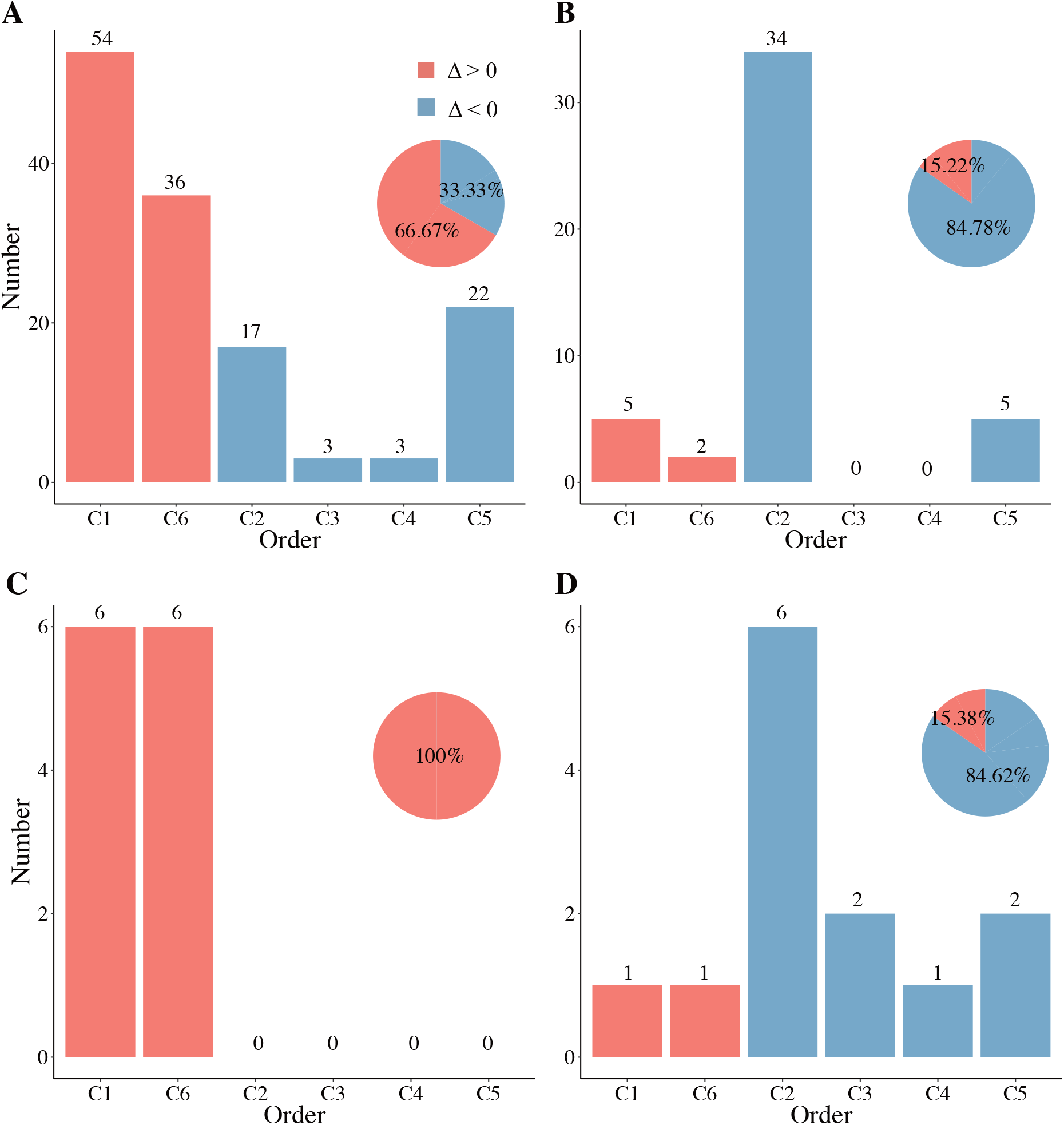
Number of significant SNPs identified by GWAS or non-parametric GWAS under different order of conditional empirical means in different datasets. The pie chart shows the percentages of significant SNPs with Δ > 0 and Δ < 0. **A-B**) Results for days to flower of cucumber from GWAS (**A**) and non-parametric GWAS (**B**). **C-D**) Results for hypoxia adaptability of chicken from GWAS (**C**) and non-parametric GWAS (**D**).

#### 2.3.2 Different methods perform diversely on the same dataset

Results in Figure 3, 4 and Figure S1 reveal another important phenomenon that different methods have diverse performance and almost all of these significant SNPs are only detected by part of these methods, which is further evidenced by manhattan plots in Figure 6A and Figure S3-S9. In general, there are some SNPs reported to be correlated with the trait of interest but only detected by a fraction of these methods (Table 4). For example, for days to flower of cucumber, there are 5 SNPs with candidate genes known to be related to this trait, but ignored by GWAS. Among these five loci, three are only detected by KS-GWAS, one by Chi-GWAS, and one by Co-GWAS (Figure 4A and Table 4). This shows the potential effect of different methods.

**Figure 6:**
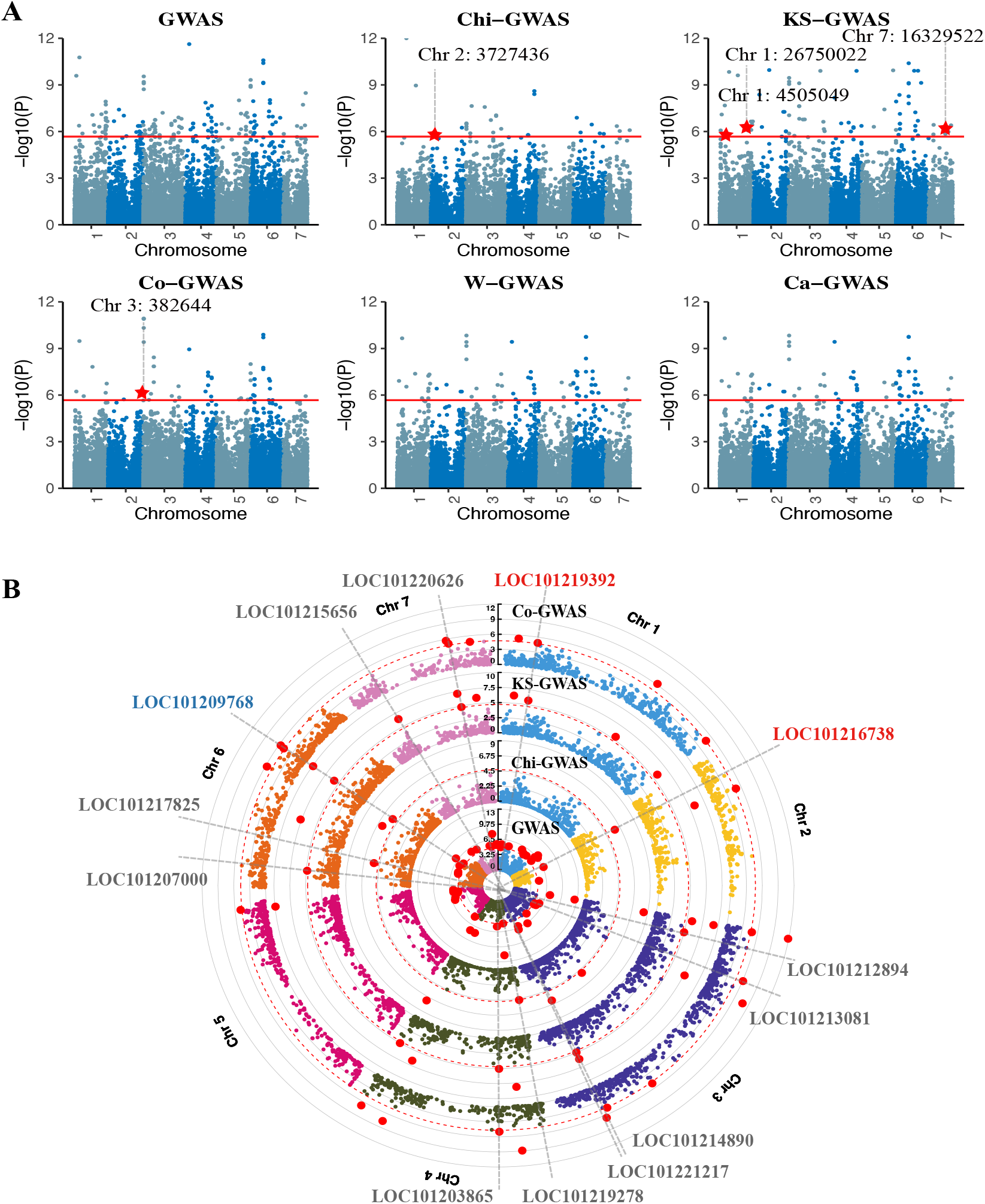
Results for days to flower of cucumber from GWAS and non-parametric GWAS. **A**) Mahanttan plot of 6 GWAS methods. The red horizontal line marks the genome-wide significance threshold adjusted by Bonferroni method. The red highlighted points stand for significant SNPs with candidate genes. **B**) Results from MAGMA analysis. In the figure, seven cucumber chromosomes (Chr 1-7) are marked by different colors, and the points represent genes in the chromosome, with significant ones colored in red. The annotated loci represent the significant genes neglected by GWAS. Among them, the blue ones were reported in the literature and red ones are also identified by non-parametric GWAS (**A**).

To further demonstrate the value of these methods, we go into details of these five candidate genes for days to flower. Among them, LOC101214996 and LOC101203410 are closely related to flowering, for instance, LOC101214996 encoding protein cup-shaped cotyledon 3 is required for the cotyledon separation process as well as the floral organ fusion (Lee et al., 2015), and LOC101203410 encoding WUSCHEL-related homeobox 13 is a potential transcription factor integrating cytokinin signaling to trigger floral induction (Li et al., 2019). Differently, LOC101219392 encoding thaumatin-like protein affects days to flower during plant growth and development (Iqbal et al., 2020). Light is known to be a key factor for flowering, therefore, genes (LOC101203816 and LOC101216738) correlated with light would exert effect on this trait as well. Specifically, Abbas and Chattopadhyay (2014) found that calmodulin-7 (LOC101203816) promotes photomorphogenic growth and light-regulated gene expression. Transcription factor PIF3 (LOC101216738) is essential for the response to various environmental signals including light and temperatures (Jiang et al., 2017).

Finally, taking the p-values from GWAS and non-parametric GWAS as input, gene association analyses are performed using MAGMA to test the correlation between genes and the trait of interest. Results in Figure 6B show that 13 significant genes are found by non-parametric methods but ignored by GWAS, where LOC101219392 and LOC101216738 are reported again, giving additional evidence to their correlations with the trait. Notably, LOC101209768 encoding tyrosine-protein phosphatase DSP1 is newly spotted. The lack of this gene would result in pleiotropic developmental defects including impaired pollen development, and thus it is related to flowering (Pu et al., 2019).

In summary, all of these methods are useful to detect part of the significant SNPs, but none of them are the best among all cases (Figure 4, 5, S1-S9). Hence, as suggested by the results shown in Simulation, we combine the power of different GWAS methods into HC-GWAS to give generally acceptable results.

#### 2.3.3 HC-GWAS spots new SNPs ignored by GWAS

Applying HC-GWAS and GWAS to seven datasets with continuous traits, new loci identified by HC-GWAS but neglected by GWAS are found in every dataset. These results manifest the advantages of HC-GWAS in the discovery of relevant genetic factors. The information of part of the significant SNPs with corresponding genes is summarized in Table 4.

Firstly, we pay attention to the genetic link to loin muscle depth (LMD) of pig (Figure 7A). LMD is an important feature of growth, affected by both nutrition and growth environment (Zhuang et al., 2019a). GWAS pinpoints 13 significant SNPs and HC-GWAS discovers 5 SNPs (Figure S1). However, 3 of these 5 SNPs are ignored by GWAS (Figure S1), and the mapped candidate gene IFNGR1 was deemed to be related to LMD, since its mRNA expression expresses differently in the muscle of adults pigs and piglets (Xing et al., 2010). Moreover, MAGMA analysis identifies a new gene (EBF1) correlated with LMD (Figure 7B), which was reported to be crucial to muscle cell maturation (Pagani et al., 2021).

**Figure 7:**
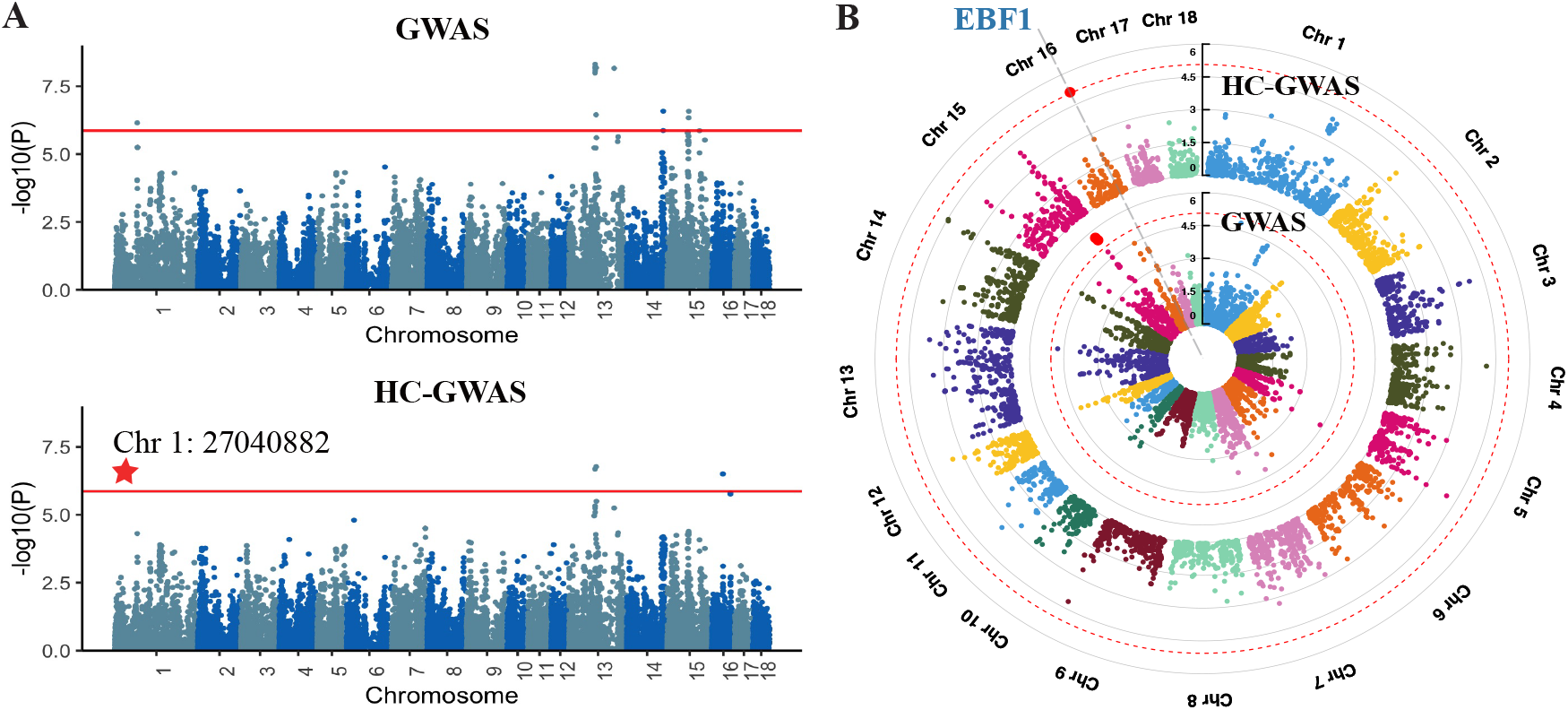
Results for loin muscle depth of pig from GWAS and HC-GWAS. **A**) Mahanttan plot of GWAS and HC-GWAS. The red horizontal line marks the genome-wide significance threshold adjusted by Bonferroni method. The red highlighted points stand for significant SNPs with candidate genes. **B**) Results from MAGMA analysis. In the figure, eighteen pig chromosomes (Chr 1-18) are marked by different colors, and the points represent genes in the chromosome, with significant ones colored in red. The annotated loci represent the significant genes neglected by GWAS. Among them, the blue ones were reported in the literature.

We next perform GWAS for the body mass index (BMI) of human. BMI is a universe standard to measure body shape and fitness. As illustrated in Figure S10, both GWAS and HC-GWAS correctly identify FTO on chromosome 16, the symbolic gene with regards to BMI, which proves their effectiveness. In addition, GWAS identifies 20 relevant SNPs, while HC-GWAS finds 45 within which 31 are new to GWAS (Figure S1). Among these new SNPs, rs58407391 resides in the gene NXPH1 that has prior known association with obesity (Oelsner et al., 2017). Thus, HC-GWAS successfully identifies this correlated gene for BMI, but GWAS fails.

For cucumber, gummy stem blight (GSB) is a serious fungal disease affecting the cultivation of cucurbitaceous vegetable crops worldwide (Stewart et al., 2015). It is caused by 3 Stagonosporopsis species, and the occurence of GSB is intensified by warm and humid environments that facilitate disease development (Gimode et al., 2021). GWAS finds 6 significant SNPs, while HC-GWAS finds 5 loci which are all new to GWAS (Figure S1). For all of these SNPs, the order of working code and that of conditional means are partially mismatched (Figure S2), demonstrating the large possibility of GWAS to lose these SNPs. Among them, the mapped gene LOC101222727, encoding small nuclear ribonucleoprotein SmD3a (Figure S10), was found to contribute to the plant immune response via the regulation of mRNA splicing of key pathogenesis factors (Golisz et al., 2021). Thus, this could be a new candidate gene for GSB resistance.

HC-GWAS also spots new SNPs ignored by GWAS for three important traits of cotton (Figure S10). Upland cotton is the most important natural fiber worldwide, and its phenotype seed index (SI) is widely used as an indicator of compost maturity, influenced by drought stress and cold environment (Chen et al., 2016; Yang et al., 2021). We perform both GWAS and non-parametric GWAS to identify significant loci, and they report exclusive SNPs and genes. GWAS identifies 277 associated SNPs, while HC-GWAS detects 37 (Figure 4C). 21 of them are only found by HC-GWAS including the gene LOC107899606. This gene encodes glutathione S-transferase U17 and was reported to play a negative role in plants drought and salt stress tolerance (Chen et al., 2012).

Boll weight (BW) is a major trait for cotton as it is key to cotton yield formation (Munir et al., 2016). Kuai et al. (2014) have discovered that sucrose transformation rate directly influences BW in cotton. GWAS finds no significant SNPs, but with the help of HC-GWAS, we identify 2 new loci (Figure S1). After careful analysis, we spot a candidate gene LOC121229363 with significant loci in its region. This gene encodes myo-inositol oxygenase and participates in various sugar metabolism and sugar sensing pathways (Gangadhar et al., 2014). Thus, this gene is very likely to be associated with BW.

Cotton fibers are developed epidermal cells of the seed coat with large amounts of cellulose and minor lignin-like components, while fiber strength (FS) well characterizes fiber qualities (Gao et al., 2019; Zhao et al., 2020). Among the 4 SNPs only detected by HC-GWAS (Figure S1), one of them has a counterpart gene LOC107915002 encoding 7-hydroxymethyl chlorophyll a reductase (HCAR). HCAR was revealed to regulate cell death and defense response against pathogen and oxidative and high light-induced damage to cells, which affects fiber length and strength (Kampire et al., 2021).

For sheep, progeny birth weight (PBW) means the weight of newborn lambs. On the SNP level, GWAS discovers 294 correlated SNPs, and HC-GWAS identifies 554 SNPs, out of which 365 SNPs are neglected by GWAS (Figure S1). On the gene level, by closely looking into the corresponding genes of each significant SNP, 4 candidate genes are found as illustrated in Figure 8A. Among them, ZEB2 was found to be influential to animals’ growth and weight traits (Martnez et al., 2017). Differently, GRIP1, TRPS1, and IGHMBP2 play a major role in syndromes with low birth weight. To be more specific, GRIP1 was indicated to influence 12q14 microdeletion syndrome, manifesting as low birth weight and developmental delay (Dria et al., 2020). Higher expression of TRPS1 produced by gestational protein restriction consequently causes low birth weight in their offspring (Assalin et al., 2019). Novel IGHMBP2 variants bring clinical diversity including a patient with SMARD1, characterized by low birth weight (Yuan et al., 2017). Furthermore, MAGMA analysis shows that there are 133 significant genes detected only by HC-GWAS including previously mentioned genes GRIP1 and IGHMBP2 (Figure 8B), which evidences the advantages of HC-GWAS. We further perform a KEGG pathway enrichment analysis of significant genes identified by HC-GWAS (Figure 8C). Among them, circadian entrainment pathway shows importance to PBW, since maternal circadian rhythm entrains fetal circadian rhythms that may subsequently affect infant outcomes (Kaur et al., 2020).

**Figure 8:**
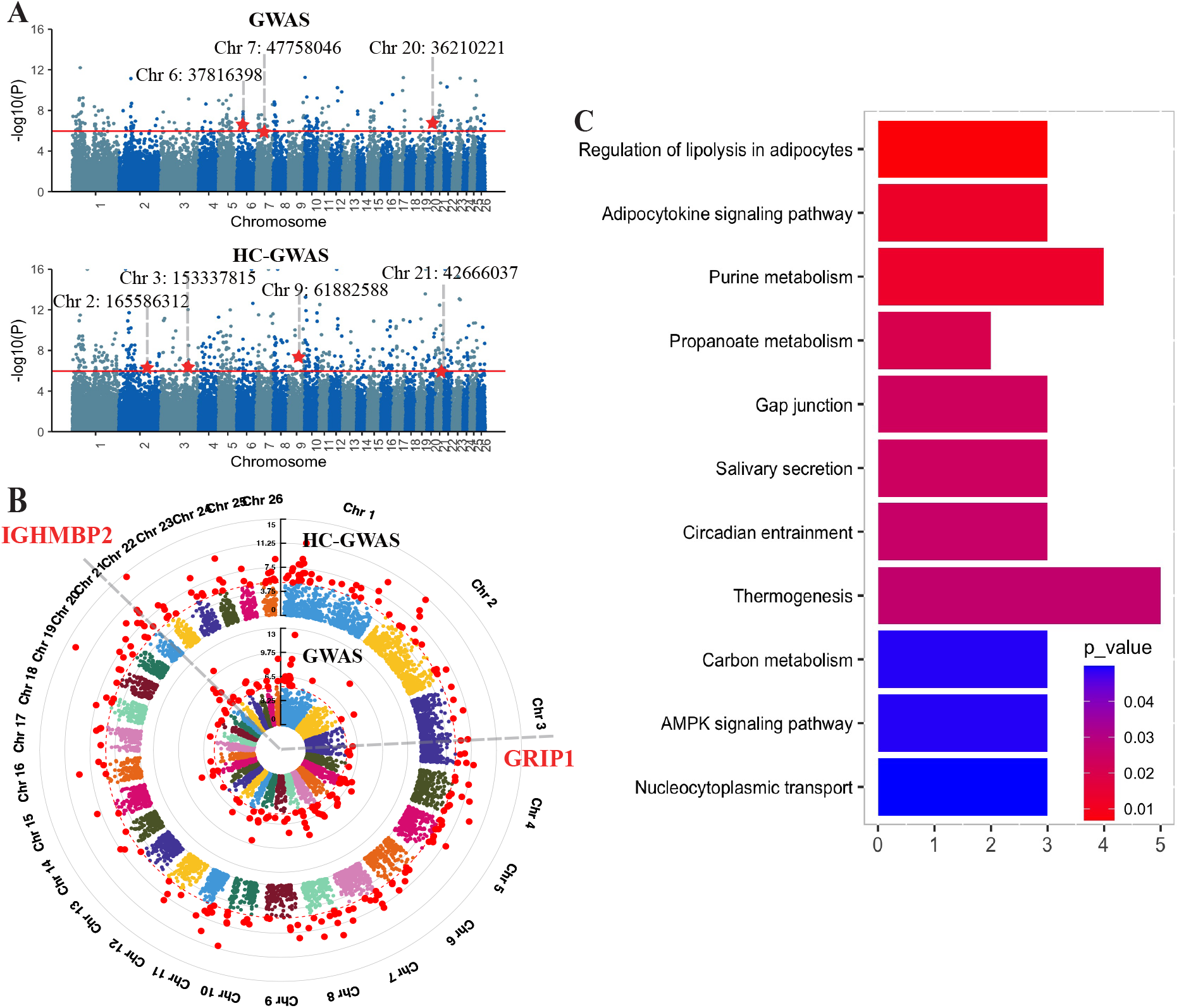
Results for progeny birth weight of sheep from GWAS and HC-GWAS. **A**) Manhattan plot of GWAS and HC-GWAS. The red horizontal line marks the genome-wide significance threshold adjusted by Bonferroni method. The red highlighted points stand for the significant SNPs with candidate genes. **B**) Results from MAGMA analysis. In the figure, twenty-six sheep chromosomes (Chr 1-26) are marked by different colors, and the points represent genes in the chromosome, with significant ones colored in red. The annotated loci represent the significant genes neglected by GWAS. Among them, red ones are also identified by non-parametric GWAS (**A**). **C**) KEGG pathway enrichment analysis for significant genes detected by HC-GWAS.

#### 2.3.4 Chi2-GWAS spotted ignored SNPs for binary traits

In this section, we present the merits of non-parametric GWAS for binary traits, Chi2-GWAS, over GWAS (based on logistic regression) in two real data sets. Firstly, we study the genetic link to abnormal bone, which refers to bone health, of Carworth Farms White (CFW) mouse. Case indicates abnormal bone, such as white and swollen bones, while control is healthy-looking bone. Both GWAS and Chi2-GWAS identify a series of related SNPs on chromosome 11 (Figure 4D and Figure 9A). Totally, GWAS finds 107 SNPs and Chi2-GWAS finds 96, but there are 15 SNPs only detected by Chi2-GWAS. Among the corresponding genes of these SNPs, Igf2bp1 is associated with bone growth and skeletal muscle development (Wang et al., 2020; Zhang et al., 2020). Moreover, MAGMA analysis illuminates two new genes RINT1 and Hoxb6 only identified by Chi2-GWAS (Figure 9B). Bi-allelic RINT1 alterations would cause a multisystem disorder including skeletal abnormalities (Cousin et al., 2019). Kappen (2016) has proved that Hoxb6 takes control of multiple independent aspects of skeletal pattern. In addition, GO pathway enrichment analysis is performed for significant genes identified by Chi2-GWAS (Figure 9C). Embryonic skeletal system morphogenesis and embryonic skeletal system development pathways are shown to be significantly correlated with bone health (Yang, 2009). Thus, the pathway enrichment analysis supports the reliability of non-parametric GWAS methods.

**Figure 9:**
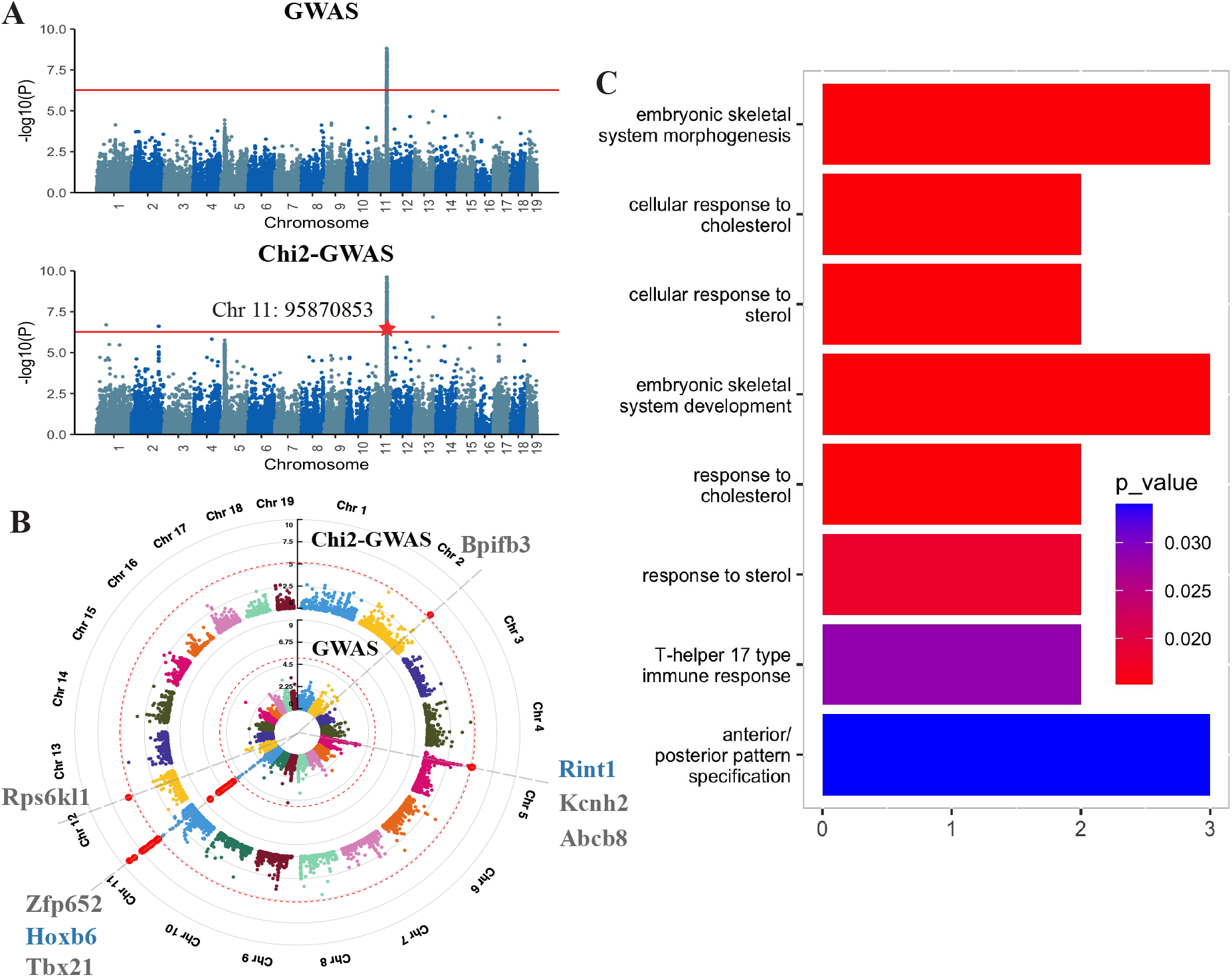
Results for abnormal bone of CFW mouse from GWAS and Chi2-GWAS. **A**) Mahanttan plot of GWAS and Chi2-GWAS. The red horizontal line marks the genome-wide significance threshold adjusted by Bonferroni method. The red highlighted points stand for significant SNPs with candidate genes. **B**) Results from MAGMA analysis. In the figure, nineteen mouse chromosomes (Chr 1-19) are marked by different colors, and the points represent genes in the chromosome, with significant ones colored in red. The annotated loci represent the significant genes neglected by GWAS. Among them, the blue ones were reported in the literature. **C**) GO pathway enrichment analysis for significant genes identified by Chi2-GWAS.

We also perform GWAS for hypoxia adaptability of Tibetan chicken who has adapted to the hypoxic and high-altitude environment for hundreds of years. Here, the hatchability under hypoxic conditions is taken as the categorical phenotype, with surviving chicks as cases and dead embryos as controls. GWAS spots 12 relevant SNPs. Chi2-GWAS discovers 20, and 13 of them are new to GWAS (Figure 4B). After close examination of the counterpart genes of these 13 SNPs, NPAS3 and EYA2 are found to be related to the trait (Figure S10). The mutants of NPAS3 homolog modulate synaptic responses in reaction to the reduction of internal oxygen levels and thus it is raised under hypoxia (Chen et al., 2019). EYA2 is regulated by epidermal growth factor receptor (EGFR) through HIF1*α*, which affects tumor microenvironment characterized by hypoxia (Liang et al., 2017).

## 3 Discussion and Conclusion

Usually, GWAS examines the effect of SNPs on the phenotype through a simple linear or logistic regression model. However, they ignore that SNPs may follow other genetic mode in practice and the coding strategy introduces unreliable numerical assumptions to SNPs. After careful theoretical analysis, we find that a partial mismatch between the order of the working code and that of the underlying true code will fail to detect true SNPs and thus cause high false negatives, which also can be explained by a general model. Here, we propose several non-parametric GWAS methods for both continuous and binary traits, and compare their power with that of GWAS in both simulated datasets and real data sets. Results show the advantages of non-parametric GWAS methods, especially in cases with partially mismatched orders between the working code and that of the conditional means of phenotype.

Despite the advantages of non-parametric methods in the scenario of partially mismatched orders, GWAS still outperforms non-parametric GWAS in other situations when the orders of working code and true code are fully matched or fully unmatched, as shown in results from simulation and real data analysis. On one hand, GWAS identifies more significant loci than non-parametric methods in nearly half of the datasets (Figrue 3), such as days to flower of cucumber, SI of cotton, and LMD of pig. On the other hand, GWAS sometimes discovers significant loci neglected by non-parametric methods. For example, for PBW of sheep, there are 3 genes, LAP3, RORA, and CDKAL1 only identified by GWAS. Importantly, they were all confirmed to be associated with lower birth weight (Everson et al., 2018; La et al., 2019; Sun et al., 2015). Similar phenomenon is also found for SI of cotton that 261 important SNPs including LOC107961898 are detected only by GWAS. LOC107961898 encoding violaxanthin de-epoxidase, chloroplastic demonstrates a positive role in both drought and salt tolerance, and thus affects SI (Sun et al., 2019), but it is ignored by HC-GWAS. All of these imply that GWAS has its advantages over non-parametric methods. These advantages may come from the fact that parametric method has a higher efficiency than non-parametric methods when the assumed model is true. To sum up, we suggest to utilize non-parametric methods as a complement instead of a substitute to GWAS to identify more associated loci. Furthermore, there is still an urgent need for better methods which can achieve high power under all circumstances, and this could be a future direction for GWAS.

## Methods

### Notations

Before presenting the methods, we introduce some notations for convenience. Let *G* be the SNP under study, *C* = {*c_i_* : *i* = 0,1, 2} be its genetic code and *Y* be the quantitative trait of interest. Usually, *c_i_* is taken as *c_i_* = *i* for *i* = 0,1,2 according to minor allele frequency, we call it default code. Denote {(*y_i_, g_i_*) : *i* = 1, 2,⋯, *n*} as *n* i.i.d samples of (*Y, G*), and define 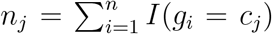 and *r_j_* = *n_j_/n* for *j* = 0,1, 2 as the number and proportion of samples whose genetic code is *c_j_*, where *I*(*A*) is the indicator function, equals 1 when *A* is true, otherwise 0. In addition, let 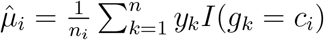 for *i* = 0,1, 2 be the conditional sample means of *Y* given *G* = *c_i_*. Finally, define 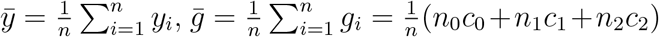 and 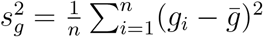.

### Analysis on linear model and general model

Typically, GWAS is based on the following linear model for quantitative trait of interest *Y*:

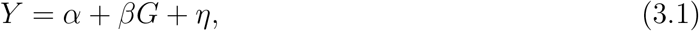

where *α* is background effect, *β* is the genetic effect, and *η* is the noise term assumed to be normal with mean 0. According to this model, finding SNPs related to the trait *Y* is to test whether *β* = 0 or not. By simple calculation, we get the least square estimator of *β* as

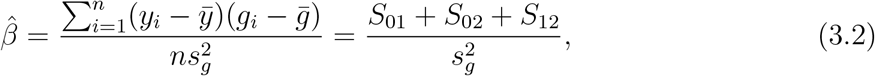

where *S_ij_* for *i,j* = 0,1, 2 is given by

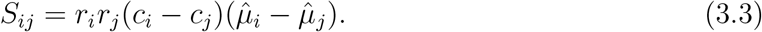

*S_ij_* can be viewed as the difference between group *i* and *j*, and the information of *β* comes from all of these difference.

Furthermore, we introduce a general model:

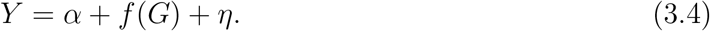

That means, the genetic effect is not directly put on the trait *Y*, but through an unknown function *f*. The linear model is a special model of this general model by taking *f*(*x*) = *βx*.

Now, we take the working code as *c_i_* = *i* for *i* = 0,1, 2, and then *β* in model (3.1) is estimated by using data from model (3.4) as,

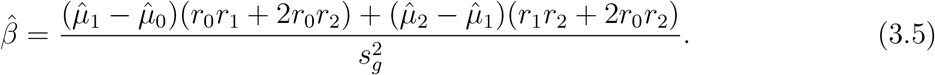

Assuming the linear model (3.1) is true, then the default code gives the order 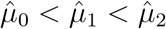, which means 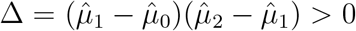, i.e, it works good for this setting. However, when this model is miss-specified, say, the SNP affect the traits via an unknown function *f* as in model (3.4), it may lead to false negatives, if we still use default code as working code. Indeed, the unknown function *f* can change the order of the default code (see Table 5). We can summarize these six cases into three types: (S1) the order of the two codes are perfectly matched (Case C1); (S2) the order of the two codes are partially mismatched (Case C2-C5); (S3) the order of the two codes are fully unmatched (Case C6).

**Table 5:** Order of mapped genetic codes compared with that of working codes

| Order of working codes: $c_0 < c_1 < c_2$ | | | | | |
| --- | --- | --- | --- | --- | --- |
| C1: $f(c_0) < f(c_1) < f(c_2)$ | C2: $f(c_1) < f(c_0) < f(c_2)$ | C3: $f(c_2) < f(c_0) < f(c_1)$ | | | |
| C4: $f(c_0) < f(c_2) < f(c_1)$ | C5: $f(c_1) < f(c_2) < f(c_0)$ | C6: $f(c_2) < f(c_1) < f(c_0)$ | | | |

For case (S1) and (S3), we all have 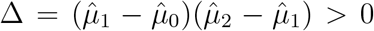. Comparing with S1, S3 simply gives an opposite conclusion on the effect of *G*, that is, a positive (negative) effect is reported mistakenly as negative (positive) effect, which is a minor problem that can be corrected by looking inside the data. However, case (S2) gives Δ < 0, which means the information from the group mean difference, 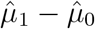 and 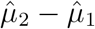, may cancel out and leads to 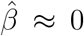 even if the true *β* is not 0, and thus reports a false negative. Typically, when *n*_1_ = *n*_2_ = *n*_3_, 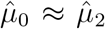 gives 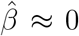, despite of 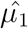. Thus, Δ < 0 is a signal for a possible false negative. In summary, when the order of working code and that of true code are partially mismatched, GWAS may have a lower power to detect meaningful SNPs.

### Non-parametric GWAS methods

To get rid of the working code of SNP, we propose the following non-parametric GWAS methods from different views of the problem.

#### Methods based on testing equality of conditional means of phenotypes given the SNP

First, we consider the case that the phenotype is continuous. On one hand, GWAS may look as the problem of testing the equality of conditional means of phenotypes given the SNP, i.e., testing the following null hypothesis

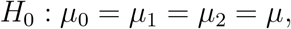

where 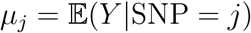, *j* = 0,1, 2.

##### W-GWAS and Ca-GWAS

It is natural to use the pairwise mean test, such as t-test and Wilcoxon rank-sum test (Datta and Satten, 2005), to compare these conditional means. Here we recommend to use Wilcoxon rank-sum test due to the fact that data are always noisy to contain some outliers. We will get three p-values from pairwise comparison, say, *p*_1_, *p*_2_ and *p*_3_. One way is to report the minimal p-value after Bonferroni correction, and the resultant method is called W-GWAS. Another way is to report a combined p-value by using the Cauchy combination (Liu and Xie, 2020), and the method is called Ca-GWAS.

##### Cauchy combination

Assume *p_i_, i* = 1, 2,⋯, *n* are p-values with any dependence structure, define

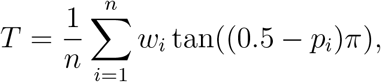

Liu and Xie (2020) show that 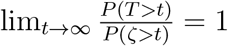, where *w_i_*’s are weights such that 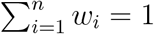 and *ζ* follows the standard Cauchy distribution. Thus, the p-value of *T*_0_, the observed value of *T*, can be estimated by *P*(*ζ* > *T*_0_).

##### Chi-GWAS

Also, we may define the test statistic for *H*_0_ as the sum of squared pairwise difference of conditional means, i.e., 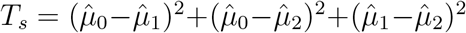, where 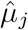, *j* = 0, 1, 2 are the estimators of *μ_j_*. This method is called Chi-GWAS. Under the null hypothesis, 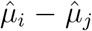 for *i* ≠ *j* are asymptotically normal distributed, thus, *T_s_* can be written in a form as 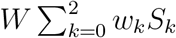, where *W* is the variance of components of *T_s_*, 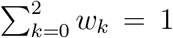, and *S_k_* is a Chi-squared random variable with degree of freedom 1. Hence, we can estimate the limit distribution of *T_s_* by 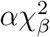, where *α* and *β* can be estimated by matching the mean and variance of 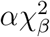 to that of *T_s_*. Details are given in supplementary material Section 1.

#### Methods based on testing equality of conditional distributions of phenotypes given the SNP

On the other hand, we may take GWAS as testing the equality of conditional distributions of phenotype *Y* given SNP, that is, testing the null hypothesis

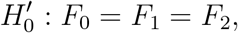

where 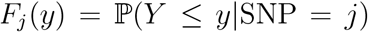, *j* = 0,1,2, are the conditional distribution of Y given SNP.

##### KS-GWAS

Kolmogorov-Smirnov (KS) (Frank and Massey, 1951) test is the first choice for comparing two distributions. Thus, our first method in this category is based on pairwise KS test, called (KS-GWAS), to report a minimal p-value after Bonferroni correction.

##### Co-GWAS

Another powerful test statistic for this problem defined by Cui et al. (2015) is

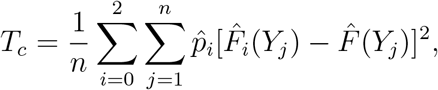

where 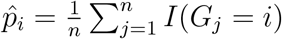 with *I*(·) being the indicator function, 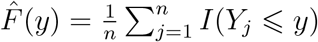, and 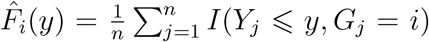. GWAS based on this statistic is called Co-GWAS. Cui et al. (2015) showed that it has a higher power on detecting associations between a categorical variable and a continuous variable, which exactly is our case. The drawback of T_c_ is that we can not get its asymptotic distribution, thus, the only way to get the p-value is permutation, which may take some time.

##### Harmonic mean combination and HC-GWAS

Wilson (2019) proposed to combine positively correlated p-values, *p*_1_, *p*_2_,⋯, *p_k_*, as 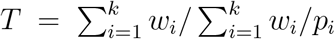, where *w_i_*’s are weights such that 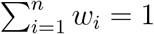. This method is shown to work better than Cauchy combination when p-values are positively correlated (Li et al., 2022). It is easy to see that p-values from different GWAS methods are positively correlated, thus, we use this method to combine power of different GWAS methods, which is called HC-GWAS. The p-value of HC-GWAS is obtained using R package “harmonicmeanp”.

#### Method for binary phenotype

Now, we consider the case where the phenotype is binary. For this case, we transform the problem of detecting the association between a SNP and the phenotype to the problem of two-sample distribution test. That is, we test the null hypothesis,

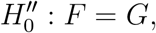

where *F* is the distribution of SNP given *Y* = 0 and *G* is the distribution of SNP given *Y* =1.

##### Chi2-GWAS

Assume that *X* = (*x*_1_,…,*x_m_*) and *Y* = (*y*_1_,…,*y_n_*) are *m* and *n* samples independently sampled from *F* and *G* respectively. To test 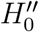, we construct an adjusted Chi-squared test statistic as 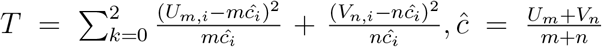, where *U_m_* = (*U*_*m*,0_, *U*_*m*,1_, *U*_*m*,2_) with 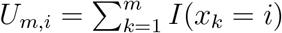 and *V_n_* = (*V*_*n*,0_, *V*_*n*,1_, *V*_*n*,2_) with 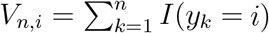, for *i* = 0,1, 2. It is shown in supplementary material Section 2 that 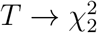 as the sample size growing to infinity. Due to the name of the statistic, we call the corresponding GWAS method as Chi2-GWAS.

### Simulation designs

In our simulation, we take the default working code as *c_i_* = *i* for *i* = 0, 1, 2, sample size as *n*_0_ = *n*_1_ = *n*_2_ = 200 and 4 different *f*(*G*) as shown below, covering all possible relationships (cases S1-S3) between working code and true code. For continuous examples, *Y* is generated from model (3.1), that is, *Y* = *f*(*G*) + *tη, η* ~ *N*(0,1), with *t* ranging from 1 to 3. The binary traits are generated by first applying the logistic transformation to *Y* in continuous examples, and then taking the transformed values above 0.5 as 1 and others as 0.

**Example A (Case S1)** *f* (*c*_0_) = 0, *f* (*c*_1_) = 1 and *f* (*c*_2_) = 2.
**Example B (Case S2)** *f* (*c*_0_) = 0, *f* (*c*_1_) = –0.3 and *f* (*c*_2_) = 0.05.
**Example C (Case S2)** *f* (*c*_0_) = 0, *f* (*c*_1_) = –0.3 and *f* (*c*_2_) = *γ*. In this example, we fix *t* = 1, and take *γ* ranging from −1 to 1.
**Example D (Case S3)** *f* (*c*_0_) = 2, *f* (*c*_1_) = 1 and *f* (*c*_2_) = 0.

### GWAS data sets

All the real data used in this study can be accessed from public websites. The human data can be obtained upon application to the UK Biobank project (Sudlow et al., 2015). Here, we randomly select 20,000 people from the UK Biobank cohort as our dataset. The cucumber data were downloaded from http://cucurbitgenomics.org/ftp/GBS_SNP/cucumber/ (Wang et al., 2018), which contains 836 cucumber accessions (Cucumis sativus L.) with 23,552 SNPs. The cotton data were downloaded from http://mascotton.njau.edu.cn/Data/GWAS_research.htm (Fang et al., 2017), containing 1,871,401 SNPs from 258 upland cotton (Gossypium hirsutum) accessions after quality control. The mouse data were downloaded from https://datadryad.org/stash/dataset/doi:10.5061/dryad.2rs41 (Parker et al., 2016), with 1150 male CFW mice (Mus musculus) phenotyped, and we filter 92,734 SNPs for GWAS. The pig data were downloaded from https://doi.org/10.6084/m9.figshare.8019551.v1 (Zhuang et al., 2019b), containing 2150 Canadian origin Duroc pig populations (Sus scrofa) with 36,740 informative SNPs. The sheep data were downloaded from https://doi.org/10.6084/m9.figshare.11859996.v1 (Esmaeili-Fard et al., 2021), comprised of 91 sheep panel (Ovis aries) with 45,342 SNPs. The chicken data were downloaded from https://www.animalgenome.org/repository/pub/CAU2018.0208/ (Jiang et al., 2018). GWAS is performed using 163 chickens (Gallus gallus), with 47,728 valid SNPs filtered after quality control.

### GWAS analysis

SNPs are filtered based on missing data rate and minor allele frequencies (MAF) with PLINK (Purcell et al., 2007). Then, imputation is performed by Beagle with default parameters (Browning et al., 2018). To avoid confounding caused by population structure and kinship in GWAS, continuous phenotypes are adjusted for the first two principal components from the PCA analysis performed by PLINK (Bani-Fatemi et al., 2016). P-values are corrected by Bonferroni method.

### MAGMA analysis

A gene-based association analysis is conducted using the Multi-marker Analysis of GenoMic Annotation (MAGMA) (de Leeuw et al., 2015), which utilizes a multiple regression method to identify multi-marker aggregated effects that account for SNP p-values and linkage disequilibrium (LD) between SNPs. The analyzed SNP set of each gene is based on whether the SNP locates in the gene body region or within extended +/− 20 kb downstream or upstream of the gene. P-values are corrected by Bonferroni method.

## Acknowledgment

This work is partially supported by Key R&D Program of Zhejiang (2021C03G2013079), and the Program of China National Tobacco Corporation (110202101032(JY-09)). In addition, we thank the participants of the included cohorts and of UK Biobank for making this work possible (UKB application 41376).

## References

N. Abbas and S. Chattopadhyay. Cam7 and hy5 genetically interact to regulate root growth and abscisic acid responses. Plant Signaling & Behavior, 9(9):e29763, 2014.

H.B. Assalin, J. Gontijo, and P.A. Boer. mirnas, target genes expression and morphological analysis on the heart in gestational protein-restricted offspring. PloS one, 14(4):e0210454, 2019.

A. Bani-Fatemi, A. Graff, C. Zai, J. Strauss, and V.D. Luca. Gwas analysis of suicide attempt in schizophrenia: main genetic effect and interaction with early life trauma. Neuroscience Letters, 622:102–106, 2016.

Y. Bossé and C.I. Amos. A decade of gwas results in lung cancer. Cancer Epidemiology, Biomarkers & Prevention: a publication of the American Association for Cancer Research, cosponsored by the American Society of Preventive Oncology, 27(4):363–379, 2018.

M. Bouaziz, C. Ambroise, and M. Guedj. Accounting for population stratification in practice: a comparison of the main strategies dedicated to genome-wide association studies. PloS one, 6(12):e28845, 2011.

B. L. Browning, Y. Zhou, and S.R. Browning. A one-penny imputed genome from nextgeneration reference panels. American journal of human genetics, 103(3):338348, 2018.

W.S. Bush and J.H. Moore. Chapter 11: Genome-wide association studies. PLoS computational biology, 8(12):e1002822, 2012.

J.H. Chen, H.W. Jiang, E.J. Hsieh, H.Y. Chen, C.T. Chien, H.L. Hsieh, and T.P. Lin. Drought and salt stress tolerance of an arabidopsis glutathione s-transferase u17 knockout mutant are attributed to the combined effect of glutathione and abscisic acid. Plant physiology, 158(1):340351, 2012.

L. T. Chen, A.Q. Sun, M. Yang, L.L. Chen, X.L. Ma, M.L. Li, and Y.P. Yin. Seed vigor evaluation based on adversity resistance index of wheat seed germination under stress conditions. The journal of applied ecology, 27(9):29682974, 2016.

P.Y. Chen, Y.W. Tsai, Y.J. Cheng, A. Giangrande, and C.T. Chien. Glial response to hypoxia in mutants of npas1/3 homolog trachealess through wg signaling to modulate synaptic bouton organization. PLoS genetics, 15(8):e1007980, 2019.

M. A. Cousin, E. Conboy, J.S. Wang, D. Lenz, T.L. Schwab, M. Williams, R.S. Abraham, S. Barnett, M. El-Youssef, R.P. Graham, L.H. Gutierrez Sanchez, L. Hasadsri, G.F. Hoffmann, N.C. Hull, R. Kopajtich, R. Kovacs-Nagy, J.Q. Li, D. Marx-Berger, V. McLin, M.A. McNiven, and E.W. Klee. Rint1 bi-allelic variations cause infantile-onset recurrent acute liver failure and skeletal abnormalities. American journal of human genetics, 105(1): 108121, 2019.

H. Cui, R. Li, and W. Zhong. Model-free feature screening for ultrahigh dimensional discriminant analysis. Journal of the American Statistical Association, 110(510):630–641, 2015.

S. Datta and G.A. Satten. Rank-sum tests for clustered data. Journal of the American Statistical Association, 100(471):908–915, 2005.

C. A. de Leeuw, J.M. Mooij, T. Heskes, and D. Posthuma. Magma: generalized gene-set analysis of gwas data. PloS Computational Biology, 11(4):e1004219, 2015.

A. Dehghan. Genome-wide association studies. In E. Evangelou, editor, Genetic Epidemiology. Methods in Molecular Biology, pages 37–49. Humana Press, New York, 2018.

S. Dria, D. Alves, M.J. Pinho, J. Pinto, and M. Leo. 12q14 microduplication: a new clinical entity reciprocal to the microdeletion syndrome? BMC medical genomics, 13(1):2, 2020.

S. M. Esmaeili-Fard, M. Gholizadeh, S.H. Hafezian, and R. Abdollahi-Arpanahi. Genes and pathways affecting sheep productivity traits: Genetic parameters, genome-wide association mapping, and pathway enrichment analysis. Frontiers in genetics, 12:710613, 2021.

T. M. Everson, T. Punshon, B.P. Jackson, K. Hao, L. Lambertini, J. Chen, M.R. Karagas, and C.J. Marsit. Cadmium-associated differential methylation throughout the placental genome: Epigenome-wide association study of two u.s. birth cohorts. Environmental health perspectives, 126(1):017010, 2018.

L. Fang, Q. Wang, Y. Hu, Y. Jia, J. Chen, B. Liu, Z. Zhang, X. Guan, S. Chen, B. Zhou, G. Mei, J. Sun, Z. Pan, S. He, S. Xiao, W. Shi, W. Gong, J. Liu, J. Ma, C. Cai, X. Zhu, W. Guo, X. Du, and T. Zhang. Genomic analyses in cotton identify signatures of selection and loci associated with fiber quality and yield traits. Nature genetics, 49(7):10891098, 2017.

J. Frank and Jr. Massey. The kolmogorov-smirnov test for goodness of fit. Journal of the American Statistical Association, 46(253):68–78, 1951.

M. Fromer, P. Roussos, S.K. Sieberts, J.S. Johnson, D.H. Kavanagh, T.M. Perumal, D.M. Ruderfer, E.C. Oh, A. Topol, H.R. Shah, L.L. Klei, R. Kramer, D. Pinto, Z.H. Gm, A.E. Cicek, K.K. Dang, A. Browne, C. Lu, L. Xie, and P. Readhead, B. Sklar. Gene expression elucidates functional impact of polygenic risk for schizophrenia. Nature neuroscience, 19 (11):14421453, 2016.

B. H. Gangadhar, J.W. Yu, K. Sajeesh, and S.W. Park. A systematic exploration of high-temperature stress-responsive genes in potato using large-scale yeast functional screening. Molecular genetics and genomics, 289(2):185201, 2014.

Z. Gao, W. Sun, J. Wang, C. Zhao, and K. Zuo. Ghbhlh18 negatively regulates fiber strength and length by enhancing lignin biosynthesis in cotton fibers. Plant science: an international journal of experimental plant biology, 286:716, 2019.

C. Giambartolomei, J. Zhenli Liu, W. Zhang, M. Hauberg, H. Shi, J. Boocock, J. Pickrell, A.E. Jaffe, CommonMind Consortium, B. Pasaniuc, and P. Roussos. A bayesian framework for multiple trait colocalization from summary association statistics. Bioinformatics, 34(15): 25382545, 2018.

W. Gimode, K. Bao, Z. Fei, and C. McGregor. Qtl associated with gummy stem blight resistance in watermelon. Theoretical and Applied Genetics, 134(2):573584, 2021.

A. Golisz, M. Krzyszton, M. Stepien, J. Dolata, J. Piotrowska, Z. Szweykowska-Kulinska, A. Jarmolowski, and J. Kufel. Arabidopsisspliceosome factor smd3 modulates immunity topseudomonas syringaeinfection. Frontiers in plant science, 12:765003, 2021.

E. Hannon, M. Weedon, N. Bray, M. O’Donovan, and J. Mill. Pleiotropic effects of trait-associated genetic variation on dna methylation: Utility for refining gwas loci. American journal of human genetics, 100(6):954959, 2017.

I. Iqbal, R.K. Tripathi, O. Wilkins, and J. Singh. Thaumatin-like protein (tlp) gene family in barley: Genome-wide exploration and expression analysis during germination. Genes, 11 (9):1080, 2020.

Y. Ji, J. Long, S.S. Kweon, D. Kang, M. Kubo, B. Park, X.O. Shu, W. Zheng, R. Tao, and B. Li. Incorporating european gwas findings improve polygenic risk prediction accuracy of breast cancer among east asians. Genetic Epidemiology, 45(5):471–484, 2021.

B. Jiang, Y. Shi, X. Zhang, X. Xin, L. Qi, H. Guo, J. Li, and S. Yang. Pif3 is a negative regulator of the cbf pathway and freezing tolerance in arabidopsis. Proceedings of the National Academy of Sciences of the United States of America, 114(32):E6695–E6702, 2017.

H. Jiang, X. Zhao, R.C.W. Ma, and X. Fan. Consistent screening procedures in highdimensional binary classification. Statistica Sinica, 32:109–130, 2022.

S.Y. Jiang, H.Y. Xu, Z.N. Shen, C.J. Zhao, and C. Wu. Genome-wide association analysis reveals novel loci for hypoxia adaptability in tibetan chicken. Animal genetics, 49(4):337339, 2018.

M.G. Kampire, R.K. Sanglou, H. Wang, B.B. Kazeem, J.L. Wu, and X. Zhang. A novel allele encoding 7-hydroxymethyl chlorophyll a reductase confers bacterial blight resistance in rice. International journal of molecular sciences, 22(14):7585, 2021.

C. Kappen. Developmental patterning as a quantitative trait: Genetic modulation of the hoxb6 mutant skeletal phenotype. PloS one, 11(1):e0146019, 2016.

S. Kaur, A.N. Teoh, N. Shukri, S.R. Shafie, N.A. Bustami, M. Takahashi, P.J. Lim, and S. Shibata. Circadian rhythm and its association with birth and infant outcomes: research protocol of a prospective cohort study. BMC pregnancy and childbirth, 20(1):96, 2020.

J. Kuai, Z. Liu, Y. Wang, Y. Meng, B. Chen, W. Zhao, Z. Zhou, and D.M. Oosterhuis. Waterlogging during flowering and boll forming stages affects sucrose metabolism in the leaves subtending the cotton boll and its relationship with boll weight. Plant science: an international journal of experimental plant biology, 223:7998, 2014.

Y. La, X. Zhang, F. Li, D. Zhang, C. Li, F. Mo, and W. Wang. Molecular characterization and expression ofspp1,lap3andlcorland their association with growth traits in sheep. Genes, 10(8):616, 2019.

B.H. Lee, J.O. Jeon, M.M. Lee, and J.H. Kim. Genetic interaction between growth-regulating factor and cup-shaped cotyledon in organ separation. Plant Signaling & Behavior, 10(2): e988071, 2015.

H. Li, H. Zhang, and H. Jiang. Combining power of different methods to detect associations in large data sets. Briefings in Bioinformatics, 23(1):1–13, 2022.

Y. Li, D. Zhang, N. An, S. Fan, X. Zuo, X. Zhang, L. Zhang, C. Gao, M. Han, and L. Xing. Transcriptomic analysis reveals the regulatory module of apple (malus domestica) floral transition in response to 6-ba. BMC Plant Biology, 19(1):93, 2019.

Y. Liang, X. Xu, T. Wang, Y. Li, W. You, J. Fu, Y. Liu, S. Jin, Q. Ji, W. Zhao, Q. Song, L. Li, T. Hong, J. Huang, Z. Lyu, and Q. Ye. The egfr/mir-338-3p/eya2 axis controls breast tumor growth and lung metastasis. Cell death & disease, 8(7):e2928, 2017.

Y. Liu and J. Xie. Cauchy combination test: a powerful test with analytic p-value calculation under arbitrary dependency structures. Journal of the American Statistical Association, 115(529):393402, 2020.

R. Martnez, D. Bejarano, Y. Gmez, R. Dasoneville, A. Jimnez, G. Even, J. Slkner, and G. Mszros. Genome-wide association study for birth, weaning and yearling weight in colombian brahman cattle. Genetics and Molecular Biology, 40(2):453459, 2017.

S. Munir, S.B. Hussain, H. Manzoor, M.K. Quereshi, M. Zubair, W. Nouman, A.N. Shehzad, S. Rasul, and S.A. Manzoor. Heterosis and correlation in interspecific and intraspecific hybrids of cotton. Genetics and molecular research, 15(2), 2016.

B. Ng, C.C. White, H.U. Klein, S.K. Sieberts, C. McCabe, E. Patrick, J. Xu, L. Yu, C. Gaiteri, D. A. Bennett, S. Mostafavi, and P.L. De-Jager. An xqtl map integrates the genetic architecture of the human brain’s transcriptome and epigenome. Nature neuroscience, 20(10):14181426, 2017.

K.T. Oelsner, Y. Guo, S.B. To, A.L. Non, and S.L. Barkin. Maternal bmi as a predictor of methylation of obesity-related genes in saliva samples from preschool-age hispanic children at-risk for obesity. BMC genomics, 18(1):57, 2017.

F. Pagani, E. Tratta, P. Dell’Era, M. Cominelli, and P.L. Poliani. Ebf1 is expressed in pericytes and contributes to pericyte cell commitment. Histochemistry and cell biology, 156 (4):333347, 2021.

C. C. Parker, S. Gopalakrishnan, P. Carbonetto, N.M. Gonzales, E. Leung, Y.J. Park, E. Aryee, J. Davis, D.A. Blizard, C.L. Ackert-Bicknell, A. Lionikas, J.K. Pritchard, and A.A. Palmer. Genome-wide association study of behavioral, physiological and gene expression traits in outbred cfw mice. Nature genetics, 48(8):919926, 2016.

J.W. Prokop, T. May, K. Strong, S.M. Bilinovich, C. Bupp, S. Rajasekaran, E.A. Worthey, and J. Lazar. Genome sequencing in the clinic: the past, present, and future of genomic medicine. Physiological Genomics, 50(8):563–579, 2018.

X. Pu, C. Meng, W. Wang, S. Yang, Y. Chen, Q. Xie, B. Yu, and Y. Liu. Dsp1 and dsp4 act synergistically in small nuclear rna 3’ end maturation and pollen growth. Plant physiology, 180(4):21422151, 2019.

S. Purcell, B. Neale, K. Todd-Brown, L. Thomas, M.A. Ferreira, D. Bender, J. Maller, P. Sklar, P.I. de Bakker, M.J. Daly, and P.C. Sham. Plink: a tool set for whole-genome association and population-based linkage analyses. American journal of human genetics, 81(3):559575, 2007.

J.E. Stewart, A.N. Turner, and M.T. Brewer. Evolutionary history and variation in host range of three stagonosporopsis species causing gummy stem blight of cucurbits. Fungal Biology, 119(5):370382, 2015.

C. Sudlow, J. Gallacher, N. Allen, V. Beral, P. Burton, J. Danesh, P. Downey, P. Elliott, J. Green, and M. Landray. Uk biobank: an open access resource for identifying the causes of a wide range of complex diseases of middle and old age. PLoS Med, 12:e1001779, 2015.

L.N. Sun, F. Wang, J.W. Wang, L.J. Sun, W.R. Gao, and X.S. Song. Overexpression of thechvdegene, encoding a violaxanthin de-epoxidase, improves tolerance to drought and salt stress in transgenic arabidopsis. 3 Biotech, 9(5):197, 2019.

X.F. Sun, X.H. Xiao, Z.X. Zhang, Y. Liu, T. Xu, X.L. Zhu, Y. Zhang, X.P. Wu, W.H. Li, H.B. Zhang, and M. Yu. Positive association between type 2 diabetes risk alleles near cdkal1 and reduced birthweight in chinese han individuals. Chinese medical journal, 128(14):18731878, 2015.

V. Tam, N. Patel, M. Turcotte, Y. Bossé, G. Paré, and D. Meyre. Benefits and limitations of genome-wide association studies. Nature reviews. Genetics, 20(8):467–484, 2019.

X. Wang, K. Bao, U.K. Reddy, Y. Bai, S.A. Hammar, C. Jiao, T.C. Wehner, A.O. Ramrez-Madera, Y. Weng, R. Grumet, and Z. Fei. The usda cucumber (cucumis sativus l.) collection: genetic diversity, population structure, genome-wide association studies, and core collection development. Horticulture research, 5:64, 2018.

Y. Wang, L. Bu, X. Cao, H. Qu, C. Zhang, J. Ren, Z. Huang, Y. Zhao, C. Luo, X. Hu, D. Shu, and N. Li. Genetic dissection of growth traits in a unique chicken advanced intercross line. Frontiers in genetics, 11:894, 2020.

D.J. Wilson. The harmonic mean p-value for combining dependent tests. Proceedings of the National Academy of Sciences of the United States of America, 116(4):11951200, 2019.

F. Xing, C. Jiang, S. Liang, L. Kang, and Y. Jiang. Genomic structure and characterization of mrna expression pattern of porcine interferon gamma receptor 1 gene. International journal of immunogenetics, 37(6):477485, 2010.

J. Yang, T. Ferreira, A.P. Morris, S.E. Medland, Genetic Investigation of ANthropometric Traits (GIANT) Consortium, DIAbetes Genetics Replication, Meta analysis (DIAGRAM) Consortium, P.A. Madden, A.C. Heath, N.G. Martin, G.W. Montgomery, M.N. Weedon, R.J. Loos, T.M. Frayling, M.I. McCarthy, J.N. Hirschhorn, M.E. Goddard, and P.M. Visscher. Conditional and joint multiple-snp analysis of gwas summary statistics identifies additional variants influencing complex traits. Nature genetics, 44(4):369S3, 2012.

Y. Yang. Skeletal morphogenesis during embryonic development. Critical reviews in eukaryotic gene expression, 19(3):197218, 2009.

Y. Yang, G. Wang, G. Li, R. Ma, Y. Kong, and J. Yuan. Selection of sensitive seeds for evaluation of compost maturity with the seed germination index. Waste management, 136: 238243, 2021.

J.H. Yuan, A. Hashiguchi, A. Yoshimura, H. Yaguchi, K. Tsuzaki, A. Ikeda, K. Wada-Isoe, M. Ando, T. Nakamura, Y. Higuchi, Y. Hiramatsu, Y. Okamoto, and H. Takashima. Clinical diversity caused by novel ighmbp2 variants. Journal of human genetics, 62(6):599604, 2017.

X. Zhang, Y. Yao, J. Han, Y. Yang, Y. Chen, Z. Tang, and F. Gao. Longitudinal epitranscriptome profiling reveals the crucial role of n6-methyladenosine methylation in porcine prenatal skeletal muscle development. Journal of Genetics and Genomics, 47(8): 466476, 2020.

W. Zhao, H. Dong, Z. Zhou, Y. Wang, and W. Hu. Potassium (k) application alleviates the negative effect of drought on cotton fiber strength by sustaining higher sucrose content and carbohydrates conversion rate. Plant physiology and biochemistry, 157:105113, 2020.

Z. Zhuang, S. Li, R. Ding, M. Yang, E. Zheng, H. Yang, T. Gu, Z. Xu, G. Cai, Z. Wu, and J. Yang. Meta-analysis of genome-wide association studies for loin muscle area and loin muscle depth in two duroc pig populations. PloS one, 14(6):e0218263, 2019a.

Z. Zhuang, S. Li, R. Ding, M. Yang, E. Zheng, H. Yang, T. Gu, Z. Xu, G. Cai, Z. Wu, and J. Yang. Meta-analysis of genome-wide association studies for loin muscle area and loin muscle depth in two duroc pig populations. PloS one, 14(6):e0218263, 2019b.

